# SOX2-phosphorylation toggles a bistable differentiation-switch in squamous cell carcinoma

**DOI:** 10.1101/2021.08.18.455699

**Authors:** Steven Hoang-Phou, Ana Sastre-Perona, Matteo Abbruzzese, Zhe Ying, Jasmin Siegle, Beatriz Aranda Orgilles, Pedro P. Rocha, Iannis Aifantis, Jane Skok, Slobodan Beronja, Markus Schober

## Abstract

The fate choice between stem cell self-renewal and differentiation is regulated by bistable transcriptional networks, which are balanced in homeostasis and imbalanced in tumors. Yet, how stem cells switch from self-renewal to differentiation remains a conundrum. Here, we discover a molecular mechanism that allows stem cell-like tumor propagating cells (TPCs) in squamous cell carcinomas (SCCs) to switch from a mutually exclusive SOX2-PITX1-TP63 self-renewal circuit to a KLF4 driven differentiation program, dependent on the relative occupancy of a novel *Klf4*-regulatory enhancer cluster (*Klf4^EC944^*) by SOX2 or KLF4, respectively. We find SOX2 occupies this site in TPCs to inhibit *Klf4* transcription, but upon phosphorylation SOX2 becomes evicted from *Klf4^EC944^*, allowing residual KLF4 to occupy this site instead, boost the expression of KLF4 and its downstream targets, and differentiate self-renewing TPCs into post-mitotic SCC cells. This mechanism allows SOX2 to promote self-renewal and tumor formation, while preserving the differentiation potential in SCC cells. Our data suggest that stochastic cell fate decisions depend on the effective concentration of enzymatically regulated transcription factors. The surprising specificity by which SOX2-phosphorylation governs the bistable *Klf4^EC944^* network-switch in SCCs reveals a conceptual framework for the identification of similar switches in other stem cell and cancer types and their potential development into cell type specific differentiation therapies for diseases in which tissue homeostasis has gone awry.

## Introduction

The binary fate choice between stem cell self-renewal or differentiation defines the rate of clonal expansion within tissues (Blanpain and Simons, 2013; Simons and Clevers, 2011). Although self-renewal and differentiation are neutrally balanced in tissue homeostasis (Klein and Simons, 2011), they become skewed towards self-renewal in cancer (Driessens et al., 2012; Klein et al., 2009). It has therefore been speculated that increased self-renewal and aberrant differentiation are the root cause of cancerous growth and the molecules controlling stemness could present effective therapeutic targets (Kreso and Dick, 2014).

Mounting evidence from genetic studies suggest that binary cell fate changes are governed by bi-stable transcriptional networks (Davis and Rebay, 2017; Young, 2011). These bi-stable networks allow cells to assume one of two mutually exclusive stable cell states, where each state is determined by master regulatory transcription factors. To maintain a specific cell state, master regulatory transcription factors co-operatively enhance each other’s expression in a positive feedback circuit, while they inhibit transcriptional programs that specify their opposing cell fate. Although cell fate defining master regulatory transcription factors and their transcriptional targets have been identified in several homeostatic tissues and cancers, it is still unclear how stem cells switch from self-renewal to differentiation.

Stratified epithelia and SCCs emerged as powerful model systems to address this long-standing question on a molecular level. Cutaneous SCCs are – similar to normal epidermis – hierarchically organized and maintained by stem cell-like cells that are located along the tumor-stroma interface (Pierce and Wallace, 1971; Schober and Fuchs, 2011). TPCs proliferate to self-renew or they differentiate into post-mitotic SCC cells without tumorigenic potential. Quantitative lineage tracing supports the idea that stem cell-like cells maintain SCC growth and their cell fate outcomes are stochastically determined when cells divide, with a bias towards self-renewal to allow for the geometric tissue expansion as squamous differentiation becomes increasingly aberrant as tumors progress (Driessens et al., 2012).

The transcription factor SOX2 is responsible for clonal expansion and SCC growth (Boumahdi et al., 2014; Siegle et al., 2014). Although *Sox2* is epigenetically repressed and not detected in normal skin epithelial cells (Arnold et al., 2011; Ezhkova et al., 2011), it becomes expressed *de novo* in TPCs (Schober and Fuchs, 2011; Siegle et al., 2014). Once SOX2 is expressed, it is specifically detected in nuclei of TPCs, but not differentiated SCC cells (Boumahdi et al., 2014; Siegle et al., 2014). SOX2 co-localizes and physically interacts with the transcription factors ΔNp63 (Jiang et al., 2020; Watanabe et al., 2014) and PITX1 (Sastre-Perona et al., 2019), and these transcription factors co-operatively promote each other’s transcription in a positive feedback circuit that drives TPC self-renewal, as they inhibit *Klf4* expression and squamous differentiation.

The transcription factor KLF4 is expressed in supra-basal epidermal keratinocytes where it promotes the differentiation of stem cell-like (basal) into post-mitotic (supra-basal) squamous epithelial cells, cornification and epidermal barrier formation (Fuchs et al., 1999). Although the regulation and function of *Klf4* in epidermal differentiation is still unclear and context dependent (Sen et al., 2012; Szigety et al., 2020), genetic gain- and loss-of-function studies suggest that KLF4 restricts SOX2, PITX1, and ΔNp63 expression and proliferation to the basal SCC layer (Sastre-Perona et al., 2019), consistent with its tumor suppressive functions (Li et al., 2012).

These gain- and loss-of-function data suggest that TPC self-renewal and differentiation are regulated by a bi-stable transcriptional network of mutually exclusive SOX2 -PITX1-ΔNp63 and KLF4 dependent gene expression programs, but it remained a conundrum how some TPCs can exit their self-renewal circuit in steady-state tumors and differentiate, when the expression of KLF4, which needs to inhibit the self-renewal circuit, is still repressed by the self-renewal program.

Here, we identify the molecular mechanism that solves this self-renewal and differentiation paradox. We find that SOX2 binds to *Klf4* enhancer cluster EC944 to inhibit *Klf4* transcription and this promotes TPC self-renewal and SCC growth. However, SOX2 is specifically evicted from EC944 when SOX2 becomes phosphorylated on Serine S39 and S253 in mouse and corresponding S37 and S251 in human SCCs allowing residual KLF4 to bind *Klf4^EC944^* instead of SOX2. KLF4 binding initiates an auto-regulatory transcriptional program promoting KLF4 expression along with other squamous differentiation markers, as increased occupancy of KLF4 at gene regulatory enhancers decommission the TPC self-renewal program. Conceptually, these data suggest that the mutually exclusive occupancy of a single transcriptional enhancer by two opposing master-regulatory transcription factors defines the binary fate choice between stem cell self-renewal and differentiation, and their occupancy of this enhancer is governed by enzymatically regulated, post-translational transcription factor modifications. The simplicity and specificity of this enzymatically regulated bi-stable switch between two stable cell states is appealing because it informs the development of differentiation therapies for cancers and other diseases where stem cell self-renewal and differentiation have gone awry.

## Results

### SOX2 is phosphorylated on Serine-37 and Serine-251 in human squamous cell carcinoma

Because we detected multiple SOX2 bands in SCC lysates (Siegle et al., 2014) and transcription factor activities can change rapidly upon phosphorylation (Whitmarsh and Davis, 2000), we wondered whether SOX2 was phosphorylated in SCCs and whether SOX2-phosphorylation could influence TPC self-renewal. To test this hypothesis, we extracted total SCC protein lysates and separated them along with calf intestinal phosphatase (CIP) treated control lysates on PhosTag gels (Figure 1A). The Phos-Tag reagent binds to phosphorylated amino acids and it reduces the mobility of phosphorylated proteins when added to poly-acrylamide gels (Kinoshita et al., 2006). Western blotting revealed four SOX2 bands and the two largest bands vanished upon CIP treatment, indicating that a fraction of SOX2 is phosphorylated in SCCs.

**Figure 1.**
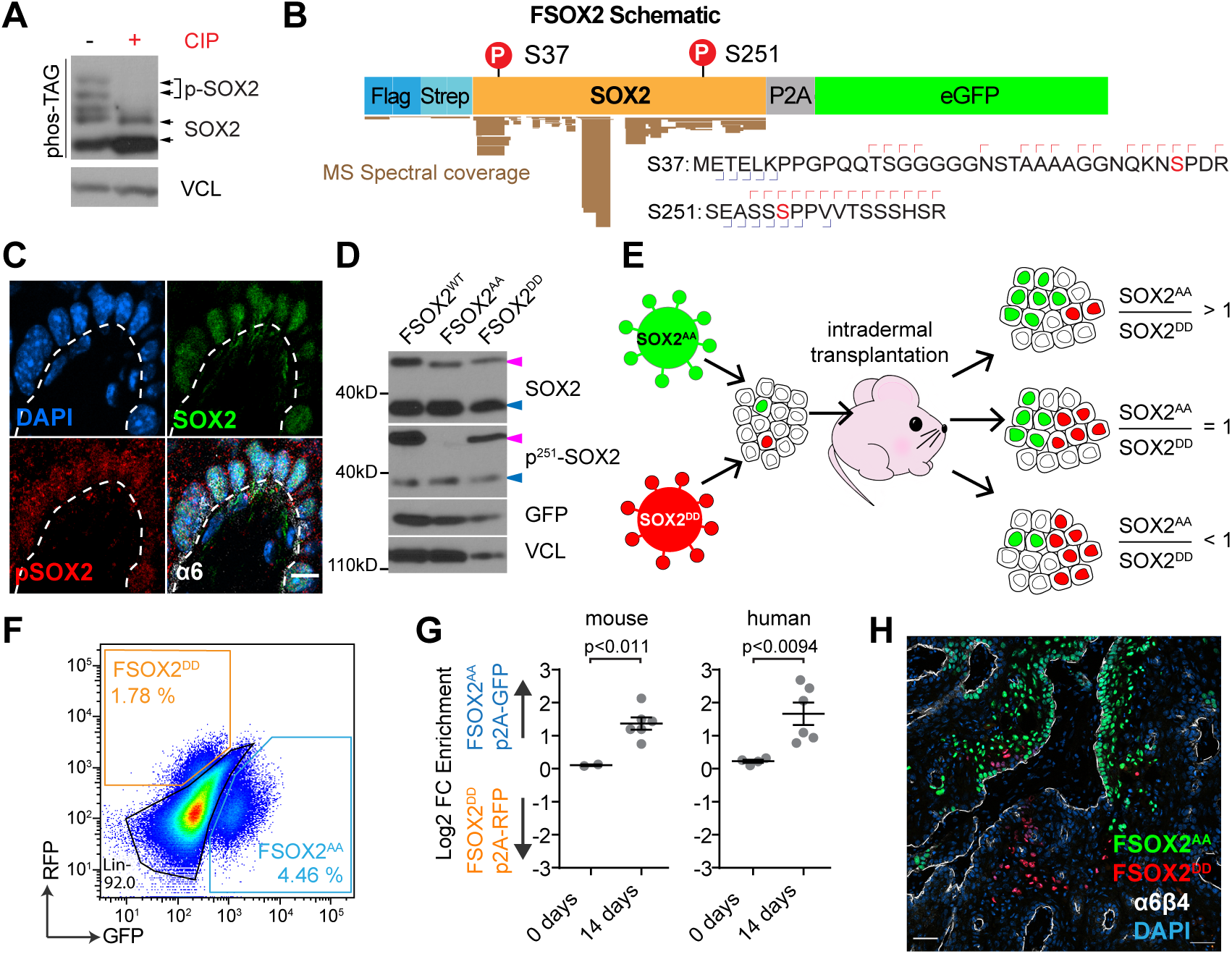
SOX2 phosphorylation reduces clonal expansion in mouse and human SCCs. **(A)** Phos-tag western blots of human SCC protein lysates treated with (+) or without (-) calf intestinal phosphatase (CIP) identifies phosphorylated and un-phosphorylated SOX2. **(B)** Schematic of human Flag-tagged SOX2 (F-SOX2), which is phosphorylated on Serine-37 and Serine-251. Brown bars indicate spectral coverage of tandem mass spectrometry results. Red and blue marks indicate fragment ion coverage. Phosphorylated Serine is shown in red. **(C)** Immunofluorescence microscopy detects SOX2 and pS251-SOX2 in nuclei of ɑ6-integrin positive SCC cells located along the tumor-stroma interface. DAPI stains nuclear chromatin. Scale bar = 10 μm. **(D)** Western blots of protein lysates from human SCC cells that express endogenous SOX2 (blue arrows) along with F-SOX2^WT^, F-SOX2^AA^, or F-SOX2^DD^ transgenes (magenta arrows). **(E)** Schematic representation of clonal competition experiment. SCC cells are infected with a pool of F-SOX2^AA^-p2A-H2B-GFP and F-SOX2^DD^-p2A-H2B-RFP containing lentiviruses at an infection rate of ∼5% before they are transplanted into the dermis where they expand into tumors. The ratio of F-SOX2^AA^-p2A-H2B-GFP over F-SOX2^DD^-p2A-H2B-RFP cells within a tumor indicates their relative expansion rates. **(F)** Flow cytometry scatter plot depicts expansion of F-SOX2^AA^-p2A-H2B-GFP and F-SOX2^DD^-p2A-H2B-RFP lineages within a representative tumor. **(G)** Scatter plot indicating Log_2_ of F-SOX2^AA^-p2A-H2B-GFP over F-SOX2^DD^-p2A-H2B-RFP ratio. n=6 independent mouse and human SCC grafts. Mean +/- s.e.m. p = unpaired parametric two tailed Student’s t-test. **(H)** Fluorescence microscopy image of murine SCC section from a clonal competition experiment showing F-SOX2^AA^-p2A-H2B-GFP enrichment along the tumor-stroma interface while F-SOX2^DD^-p2A-H2B-RFP cells are detected in supra-basal and differentiated areas. ɑ6-integrin marks tumor-stroma interface. DAPI identifies nuclear chromatin. Scale bar = 50 μm

To determine which SOX2 amino acids are phosphorylated in SCCs, we first fused two FLAG and two Streptavidin tags to the N-terminal end of human SOX2 (*F-SOX2*) (Figure 1B). Human and mouse SOX2 are almost identical in their amino acid sequences and they differ only by an insertion of two glycine residues at position 23-24 in mouse SOX2. We expressed our F-SOX2 transgenes in human SCC cultures close to endogenous levels, purified large amounts of F-SOX2 by immuno-precipitation, digested it with cyanogen bromide-Trypsin, and identified phosphorylated peptides with tandem mass-spectrometry (**Figure S1A-C**). This approach identified two phosphorylated peptides with fragmentation patterns corresponding to human SOX2 that is phosphorylated on Serine 37 (S37) and Serine 251 (S251) (Figure 1B**, S1D-E**). Confocal microscopy with site specific phospho-SOX2 antibodies detected pS253-SOX2 in mouse (Figure 1C) and corresponding pS251-SOX2 in human **(Figure S1F**) SCC sections, indicating that endogenously expressed SOX2 is phosphorylated in primary SCCs.

To functionally test if SOX2 S37 and S251 phosphorylation affects SOX2 activity, we mutated both serine residues in our F-SOX2 transgene to either alanine (A) or aspartic acid (D) before we expressed them in SCC cells. Consistent with the notion that alanine cannot be phosphorylated and aspartic acid is similar to phosphorylated serine in charge and structure, we detected endogenously expressed SOX2 with anti-pS251-SOX2 antibodies along with the larger F-SOX2^WT^ and F-SOX2^DD^, but not the F-SOX2^AA^ transgene by western blotting (Fig 1D). In addition, anti-FLAG antibodies detected only a single F-SOX2^AA^ band on Phos-Tag gels, whereas F-SOX2^WT^ showed a multi-band pattern (**Figure S1G**), mirroring that of endogenously expressed SOX2 (Figure1A). Together, these data show that endogenous SOX2 and F-SOX2 transgenes are phosphorylated on S37 and S251 in SCCs and these two sites are either the only phosphorylated SOX2 amino acids or their phosphorylation is required for additional phosphorylation events in SCC cells.

### F-SOX2^AA^ clones outcompete F-SOX2^DD^ expressing clones in squamous cell carcinomas

To test if SOX2-phosphorylation affects its activity, we turned to a clonal competition model (Ge et al., 2017; Siegle et al., 2014), which allows us to directly compare the relative expansion of *F-SOX2^DD^;H2B-RFP* and *F-SOX2^AA^;H2B-GFP* expressing cells within the same tumor over time. We generated lentiviruses for each transgene and combined them at equivalent titers before infecting primary TPC cultures at low multiplicity of infection. FACS analyses confirmed ∼5% of TPCs expressed either *F-SOX2^DD^;H2B-RFP* or *F-SOX2^AA^;H2B-GFP* before we transplanted them orthotopically into mouse dermis (**Figure S1H**). Two weeks after transplantation, we measured how *F-SOX2^AA^;H2B-GFP* cells expanded in comparison to *F-SOX2^DD^;H2B-RFP* cells within the same tumor (Figure 1E). FACS analyses revealed that SOX2^AA^ expressing clones expand faster than SOX2^DD^ expressing clones in both, mouse and human SCC transplantation models (Figure 1F-G**, S1H**). Likewise, confocal microscopy also detected more *F-SOX2^AA^;H2B-GFP* compared to *F-SOX2^DD^;H2B-RFP* expressing cells in SCC sections (Figure 1H). Interestingly, *F-SOX2^AA^;H2B-GFP* expressing cells were frequently detected in the basal SCC layer along the tumor-stroma interface, whereas *F-SOX2^DD^;H2B-RFP* expressing cells were more commonly detected in differentiated, post-mitotic tumor parts. This differential localization pattern suggested that SOX2 phosphorylation inhibits TPC self-renewal or enables the differentiation of TPCs into post-mitotic SCC cells even when endogenous SOX2 is also expressed in these SCC cells.

### Ectopic SOX2 expression accelerates squamous cell carcinoma initiation in a phosphorylation dependent manner

Because *Sox2* is epigenetically repressed and not detected in normal skin epithelial cells (Arnold et al., 2011; Ezhkova et al., 2011), but becomes *de novo* expressed in TPCs of cutaneous SCCs (Boumahdi et al., 2014; Schober and Fuchs, 2011; Siegle et al., 2014), we wondered whether ectopic SOX2 expression in skin epithelial cells would promote SCC initiation and growth and whether tumor initiation rates would vary with SOX2^WT^, SOX2^AA^ and SOX2^DD^ expression. To address these questions, we transduced *Hras^G12V-KI^;R26-LSL-YFP* mice at e9.5 (Beronja et al., 2010) with lentiviruses expressing either CRE alone (red), or CRE along with SOX2^WT^ (brown), SOX2^AA^ (blue), or SOX2^DD^ (orange) and measured their relative tumor initiation rates over time (Figure 2A). CRE expression activated oncogenic *Hras^G12V^* along with YFP expression in transduced cells and mice with transduced epidermis were born asymptomatic and they developed normally after birth. However, some *Hras^G12V^* expressing mice began to develop tumors after 40+ days (Figure 2B). Ectopic F-SOX2^WT^ or F-SOX2^AA^ expression significantly enhanced the rate of tumor initiation in this HRAS^G12V^ driven carcinogenesis model. However, this accelerated tumor initiation rate was significantly reduced in F-SOX2^DD^ expressing mice. Similarly, ectopic *Hras^G12V^* expression in FACS isolated interfollicular (ɑ6^hi^, Sca1^hi^) skin epithelial cells (**Figure S2A**), resulted in the development of some slow growing and often spontaneously regressing tumors after intra-dermal transplantation. However, if we co-transduced these cells with F-SOX2^WT^ or F-SOX2^AA^ we measured robust and significantly accelerated tumor growth, whereas HRAS^G12V^;F-SOX2^DD^ expressing tumors expanded significantly slower (Figure 2C**, S2B**). These data suggest ectopic SOX2 expression promotes HRAS^G12V^ driven squamous carcinogenesis in skin, and this activity is reduced in phosphorylation mimetic *SOX2^DD^* mutants.

**Figure 2.**
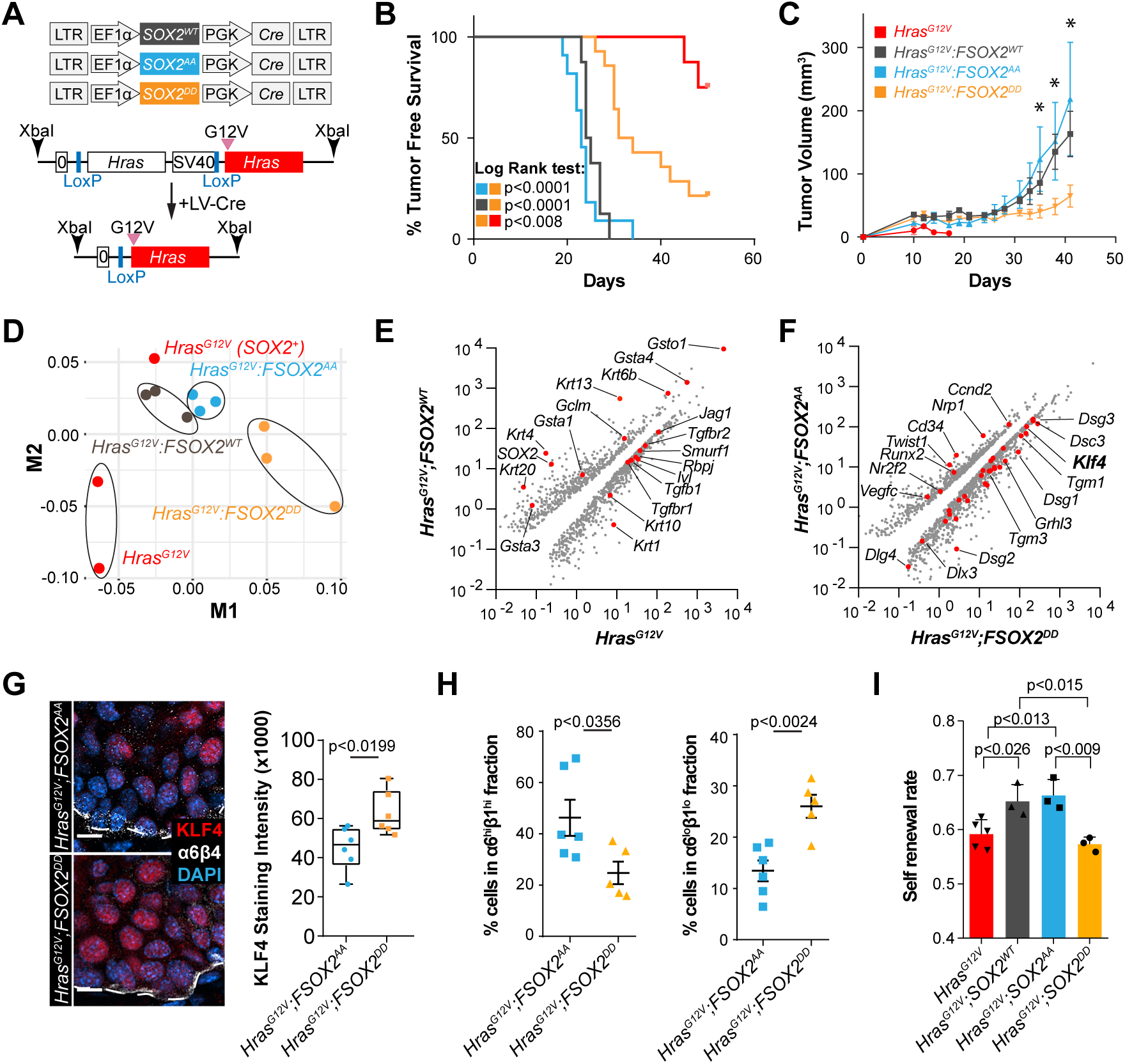
Ectopic SOX2 expression enhances self-renewal and tumor initiation in a *Hras^G12V^* driven skin cancer model. **(A)** *Hras^G12V^* knock-in mouse embryos are infected in utero with lentiviruses that express either *SOX2^WT^*, *SOX2^AA^*, or *SOX2^DD^* along with CRE recombinase at e9.5. **(B)** Kaplan Meier graphs showing tumor free survival times. p<0.0001 for all comparisons except Hras^G12V^ vs *Hras^G12V^;F-SOX2^DD^* p = 0.0075 and *Hras^G12V^;F-SOX2^WT^* vs *Hras^G12V^;F-SOX2^AA^* p = 0.2354. p = Log-rank (Mantel-Cox) test. **(C)** Tumor growth analyses of *Hras^G12V^* (n=21), *Hras^G12V^;F-SOX2^WT^* (n=15), *Hras^G12V^;F-SOX2^AA^* (n=15), *Hras^G12V^;F-SOX2^DD^* (n=15). Mean +/- s.e.m ; n=3 independent experiments; all marked * at least p<0.002, p = two-tailed t-test with Holm-Sidak correction. **(D)** Principle component analysis (PCA) of RNA-seq data from *Hras^G12V^*, *Hras^G12V^;F-SOX2^WT^*, *Hras^G12V^;F-SOX2^AA^*, *Hras^G12V^;F-SOX2^DD^* tumors. Note: One Hras^G12V^ tumor expressed endogenous *Sox2 de novo* and is more closely related to *Hras^G12V^;F-SOX2^WT^* and *Hras^G12V^;F-SOX2^AA^* tumors. **(E-F)** Scatter plots depicting differentially expressed genes (q<0.05) in *Hras^G12V^* vs. *Hras^G12V^;F-SOX2^WT^* **(E)**, or *Hras^G12V^;F-SOX2^AA^* vs *Hras^G12V^;F-SOX2^DD^* tumors **(F)**. Note: the *Hras^G12V^;SOX2^+^* tumor was excluded from the *Hras^G12V^* data set **(E)**. Selected differentiation and self-renewal markers are indicated in red. **(G)** Representative confocal microscopy images of KLF4 stained *Hras^G12V^;F-SOX2^AA^* and *Hras^G12V^;F-SOX2^DD^* tumors. Box-plots depicting KLF4 staining intensities in these tumors. n=6. P = unpaired two-tailed parametric Student’s t-test. Scale bar = 10 µm. **(H)** Relative fraction of ɑ6^hi^/β1^hi^ and ɑ6^lo^/β1^lo^ cells in GFP-labelled *Hras^G12V^;F-SOX2^AA^* or *Hras^G12V^;F-SOX2^DD^* SCC-lineage. Mean +/- s.e.m; p = unpaired two-tailed parametric Student’s t-test. **(I)** Bar graph indicating self-renewal rate in *Hras^G12V^*, *Hras^G12V^;F-SOX2^WT^*, *Hras^G12V^;F-SOX2^AA^*, *Hras^G12V^;F-SOX2^DD^* tumors. n > 3 tumors. Mean +/- s.e.m. p = unpaired two tailed parametric Student’s t-test.

### *De novo* SOX2 expression inhibits squamous differentiation gene expression

Although differential gene expression profiles between DMBA initiated control and *Sox2* knock-out SCCs identified a SOX2 regulated gene set in established SCCs (Boumahdi et al., 2014), it is still unclear how *de novo* SOX2 expression promotes SCC initiation and whether and how SOX2 phosphorylation influences the process. Therefore, we isolated lineage marked cells from *Hras^G12V^*, *Hras^G12V^;F-SOX2^WT^*, *Hras^G12V^;F-SOX2^AA^*, and *Hras^G12V^;F-SOX2^DD^* expressing tumors and determined their gene expression profiles by RNA-seq (**Figure S2C**). Principle component analyses revealed that *Hras^G12V^;F-SOX2^WT^* and *Hras^G12V^;F-SOX2^AA^* expressing tumors are closely related to one another and they are distinct from *Hras^G12V^* as well as *Hras^G12V^;F-SOX2^DD^* expressing tumors (Figure 2D) consistent with their tumor growth data (Figure 2C). Intriguingly, one *Hras^G12V^* tumor was closely related to *Hras^G12V^;F-SOX2^WT^* and *Hras^G12V^;F-SOX2^AA^* tumors, and closer data inspection revealed that it expressed SOX2 *de novo,* while SOX2 was not detected in the other two *Hras^G12V^* tumors (**Figure S2D**). Confocal immunofluorescence microscopy confirmed SOX2 expression and phosphorylation in this *Hras^G12V^* tumor (**Figure S2E**). This serendipitous finding validates our ectopic SOX2 transgene expression approach as pathologically relevant and powerful to dissect early stages of SCC initiation and promotion.

Differential gene expression analyses between *Hras^G12V^* and *Hras^G12V^;F-SOX2^WT^* driven tumors revealed SOX2 inhibits the tumor suppressive NOTCH (*Jag1*, *Rbpj*) (Alcolea et al., 2014; Fortunel et al., 2003; Rangarajan et al., 2001; South et al., 2014) and TGFβ (*Tgfb1*, *Tgfbr1*, *Tgfbr2*, *Smurf1*) (Guasch et al., 2007; White et al., 2010) signaling pathways along with spinous differentiation (*Krt1* and *Krt10*) markers (Figure 2E). In addition, we uncovered a significant increase in TPC marker expression (*Cd34* (Malanchi et al., 2008), *Vegfc* (Beck et al., 2011; Lichtenberger et al., 2010), *Nrp1* (Beck et al., 2011; Siegle et al., 2014), and *Twist1* (Beck et al., 2015)) and suppression of squamous differentiation genes (*Klf4* (Fuchs et al., 1999; Li et al., 2019), *Grhl3* (Yu et al., 2006), *Dsg1*, *Dsg2*, *Dsc1*, *Dsc2* (Dusek et al., 2007), *Dlx3* (Morasso et al., 1996; Palazzo et al., 2016) and *Tgm1*, *Tgm3* (Eckert et al., 2005)) in *Hras^G12V^;F-SOX2^AA^* compared to *Hras^G12V^;F-SOX2^DD^* SCCs (Figure 2F). These data suggest SOX2 promotes tumor initiation by suppressing squamous differentiation and this activity is inhibited by SOX2 phosphorylation.

### SOX2-phosphorylation inhibits progenitor cell self-renewal, SCC initiation and growth

Consistent with the idea that SOX2 phosphorylation inhibits TPC self-renewal and promotes their differentiation, we detected higher KLF4 expression in *Hras^G12V^;F-SOX2^DD^* compared to *Hras^G12V^;F-SOX2^AA^* SCCs (Figure 2G). In addition, FACS analyses revealed a significantly reduced fraction of undifferentiated (ɑ6^hi^β1^hi^) SCC cells alongside a larger cohort of differentiated (ɑ6^lo^β1^lo^) SCC cells in *Hras^G12V^;F-SOX2^DD^* tumors (Figure 2H**, S2C**). To measure TPC self-renewal rates, we also labeled cycling cells with EdU (5-Ethynyl-2’deoxyuridine) for 2 hrs, followed by an 8 hr BrdU (5-Bromo-2’deoxyuridine) pulse (**Figure S2F**). Next, we determined the fraction of undifferentiated cells that stained positive for EdU but not KRT10, over all EdU labelled cells, including KRT10 positive, differentiated SCC cells with confocal microscopy (Ying et al., 2018). These experiments also suggested TPC self-renewal is significantly reduced in *Hras^G12V^;F-SOX2^DD^* compared to *Hras^G12V^;F-SOX2^AA^* SCCs (Figure 2I) and SOX2 phosphorylation inhibits TPC self-renewal by accelerating their differentiation into post-mitotic SCC cells.

### SOX2-phosphorylation allows TPCs to differentiate in established SCCs

To test if SOX2 phosphorylation also effects SCC maintenance, we turned to dimethylbenzanthracene (DMBA) initiated SCCs, which mirror the genetic heterogeneity of human SCCs (Nassar et al., 2015) and are *Sox2* dependent (Boumahdi et al., 2014; Siegle et al., 2014). We isolated TPCs from these tumors, transduced them with either *F-SOX2^WT^, F-SOX2^AA^,* or *F-SOX2^DD^* transgenes before we deleted endogenous *Sox2* with CRISPR-Cas9 (Figure 3A). Next, we selected single cell clones and verified *Sox2* deletion and F-SOX2 transgene expression (Figure 3B). Western blotting with SOX2 antibodies detected F-SOX2^WT^, F-SOX2^AA^, or F-SOX2^DD^ transgene expression, but no endogenously expressed SOX2. Furthermore, pS251-SOX2 antibodies identified F-SOX2^WT^ and F-SOX2^DD^, but not F-SOX2^AA^ in these TPC clones. We recovered *Sox2^KO^;F-SOX2^AA^* clones at a higher frequency compared to *Sox2^KO^;F-SOX2^WT^* or *Sox2^KO^;F-SOX2^DD^* clones consistent with the idea that unphosphorylated SOX2 is more active in TPCs (**Figure S3A**).

**Figure 3.**
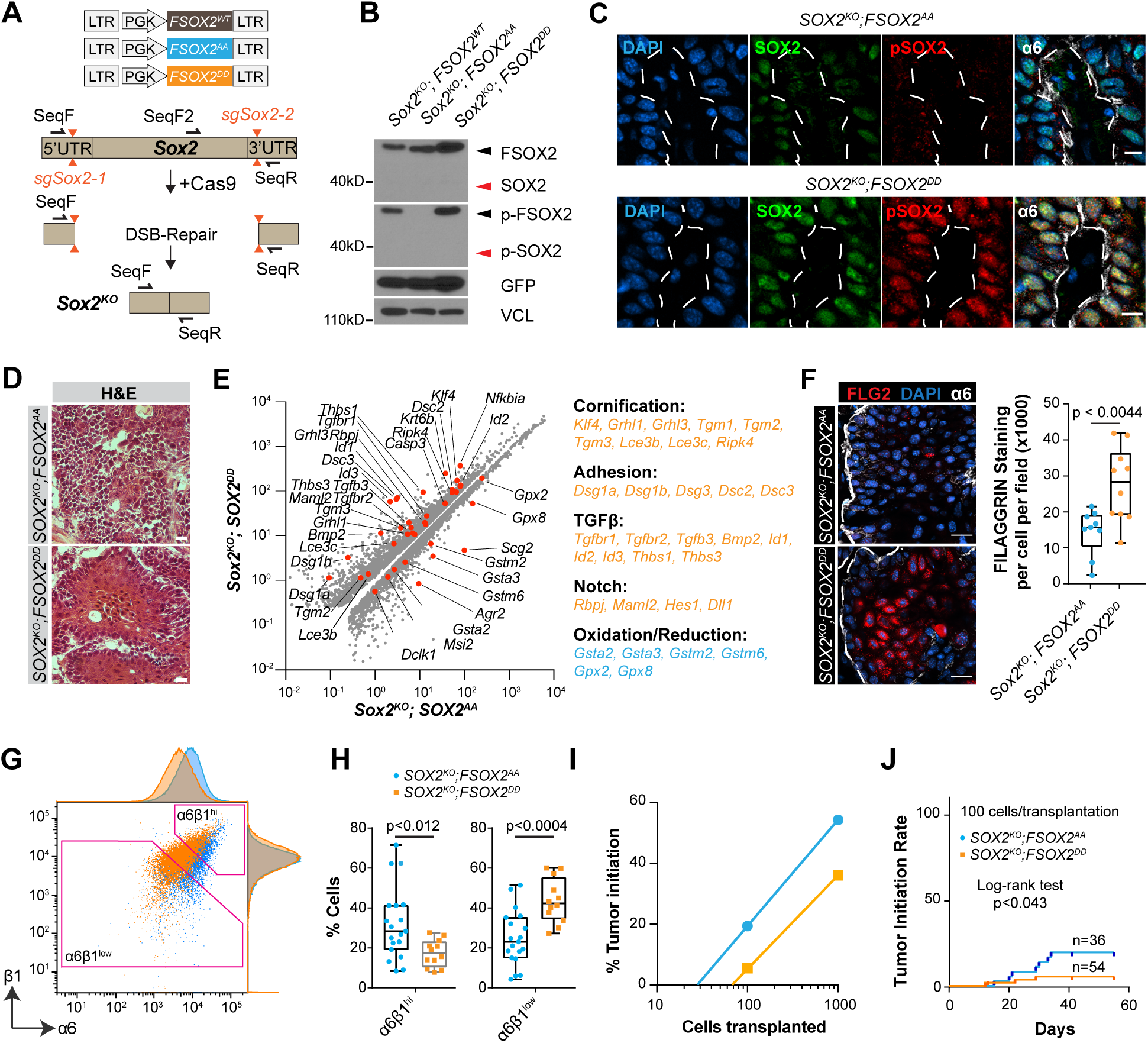
SOX2 phosphorylation enhances differentiation in SCCs. **(A)** *Sox2^KO^* and SOX2-mutant replacement strategy. SOX2 function is essential in SCC cells. Therefore, we expressed F-SOX2^WT^, F-SOX2^AA^, or F-SOX2^DD^ transgenes before we targeted the 5’ and 3’ UTRs with sgRNA’s to delete endogenous *Sox2* with CRISPR/Cas9. **(B)** SOX2 and pS251-SOX2 western blotting confirms effective *Sox2* deletion and F-SOX2^WT^, F-SOX2^AA^, or F-SOX2^DD^ transgene expression. **(C)** Confocal micrographs of SOX2 and pS251-SOX2 stained *Sox2^KO^;F-SOX2^AA^* and *Sox2^KO^;F-SOX2 ^DD^* SCC sections. **(D)** Hematoxylin & Eosin staining reveals a more disorganized architecture in *Sox2^KO^;F-SOX2^AA^* compared to *Sox2^KO^;F-SOX2^DD^* SCC sections. Scale bars = 25µm. **(E)** Scatter plot depicting differentially expressed genes (q<0.05) between *Sox2^KO^;F-SOX2^AA^* vs *Sox2^KO^;F-SOX2^DD^* SCCs. Select differentiation and SCC marker genes are depicted in red. Table highlights gene sets enriched in *Sox2^KO^;F-SOX2^DD^* (orange) and *Sox2^KO^;F-SOX2^AA^* (blue) SCCs. **(F)** Confocal microscopy section reveals increased FLG2 staining in *Sox2^KO^;F-SOX2^DD^* compared to *Sox2^KO^;F-SOX2^AA^* SCCs. Box-plots illustrate differences in FLG2 staining intensities. n=3 tumors with 3 fields per tumor. p = unpaired parametric two-tailed t-test. **(G-H)** Representative scatter plots with adjunct histograms illustrating ɑ6- and β1-integrin staining intensities in *Sox2^KO^;F-SOX2^AA^* (blue) and *Sox2^KO^;F-SOX2^DD^* (orange) SCCs **(G)**. Relative fraction of ɑ6^hi^/β1^hi^ and ɑ6^lo^/β1^lo^ cells in GFP-labelled *Sox2^KO^;F-SOX2^AA^* and *Sox2^KO^;F-SOX2^DD^* SCC-lineage **(H)**. n = 19 *Sox2^KO^;F-SOX2^AA^*. n = 12 *Sox2^KO^;F-SOX2^DD^*. p = unpaired parametric two tailed Student’s t-test. **(I-J)** Calculated tumor initiation frequency **(I)** and tumor initiation rate **(J)** after serial transplantation of lineage marked *SOX2^KO^;F-SOX2^AA^* or *Sox2^KO^;F-SOX2^DD^* cells. p = Mantel-Cox Log-rank test.

When we transplanted these clones into mouse dermis, they formed SCCs in which SOX2 was either unphosphorylated or constitutive SOX2 phosphorylation was simulated (Figure 3C). We noticed *F-SOX2^DD^* was consistently higher expressed than *F-SOX2^AA^* in our rescued *Sox2^KO^* clones. This expression difference suggests phosphorylated SOX2 is less active but increased expression might partially compensate for this activity loss. Nevertheless, *Sox2^KO^;F-SOX2^DD^* SCCs had a more differentiated appearance than *Sox2^KO^;F-SOX2^AA^* tumors (Figure 3D) and their differential gene expression profiles featuring increased cornification, adhesion, TGFβ- and NOTCH-activation along with reduced Oxidation/Reduction (Figure 3E) similar to what we observed in our tumor initiation model (Figure 2F).

This increase in squamous differentiation and cornification was also reflected in a significantly higher FLG2 expression (Figure 3F) along with a reduced fraction of undifferentiated (ɑ6^hi^/β1^hi^) and an increased fraction of differentiated (ɑ6^lo^/β1^lo^) SCC cells in *Sox2^KO^;F-SOX2^DD^* compared to *Sox2^KO^;F-SOX2^AA^* SCCs (Figure 3G-H**, S3B**). To functionally test if these changes in tumor composition and differentiation affect the tumorigenic SCC cell fraction, we isolated an equal number of lineage-marked *Sox2^KO^;F-SOX2^AA^* and *Sox2^KO^;F-SOX2^DD^* cells and re-transplanted them in a limited dilution series into mouse dermis. *Sox2^KO^;F-SOX2^DD^* SCC cell transplants initiated tumors at a lower frequency (Figure 3I) and a significantly slower rate (Figure 3J) when compared to *Sox2^KO^;F-SOX2^AA^* SCCs. These data suggest phosphorylation constrains SOX2 activity, resulting in reduced TPC self-renewal and SCC growth due to increased differentiation, but it doesn’t completely inhibit SOX2 function.

### Un-phosphorylated and phospho-mimetic SOX2 show similar chromatin binding patterns

To investigate how SOX2-phosphorylation inhibits TPC self-renewal, we prepared nuclear chromatin from *Hras^G12V^;F-SOX2^WT^*, *Hras^G12V^;F-SOX2^AA^*, and *Hras^G12V^;F-SOX2^DD^*, as well as *Sox2^KO^;F-SOX2^WT^, Sox2^KO^;F-SOX2^AA^, and Sox2^KO^;F-SOX2^DD^* SCC models for chromatin immunoprecipitation sequencing (ChIP-seq). To accurately normalize these ChIP-seq data, we spiked equal amounts of chromatin we isolated from a human SCC cell line into each murine ChIP-seq sample before immuno-precipitation. After each sample was sequenced, we aligned mouse and human sequences that co-precipitated with SOX2 against the respective reference genomes and normalized mouse relative to their corresponding human sequence counts. Although the DNA-binding ability of transcription factors is often inhibited when they become phosphorylated, we found F-SOX2^WT^, F-SOX2^AA^ and F-SOX2^DD^ ChIP-seq profiles were highly concordant and more than 80% of SOX2-peaks contained the SOX-family motif at their peak summits (Figure 4A-B). These data suggest our ChIP-seq results are specific and SOX2-phosphorylation does not globally interfere with its DNA-binding ability.

**Figure 4.**
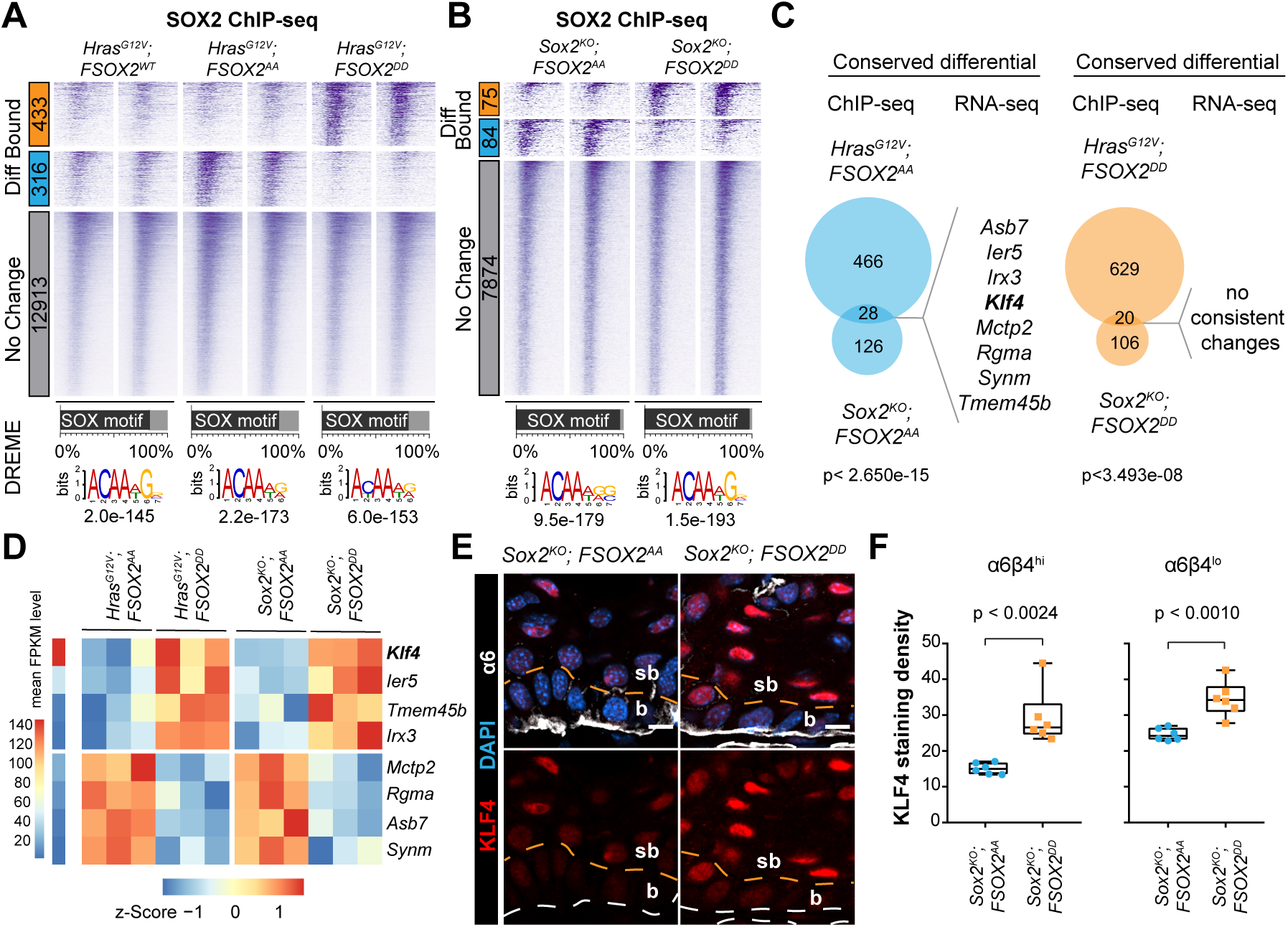
SOX2-phosphorylation affects *Klf4* in a small subset of SOX2-bound sites. **(A-B)** Heatmap representation of SOX2 bound gene regulatory elements that are significantly enriched (fold change > 1.5, p < 0.05) in *Hras^G12V^;FSOX2^AA^* or *Hras^G12V^;FSOX2^DD^* **(A)** and *Sox2^KO^;F-SOX2^AA^* or *Sox2^KO^;F-SOX2^DD^* **(B)** SCCs. Most SOX2 bound gene regulatory elements are unaffected by SOX2-phosphorylation. DREME discovered SOX-like motif and percentage of peaks containing the SOX motif. n=2 per genotype. **(C)** Intersection of differentially bound and expressed genes in *Hras^G12V^;FSOX2^DD^* and *Sox2^KO^;F-SOX2^DD^* or *Hras^G12V^;FSOX2^AA^* and *Sox2^KO^;F-SOX2^AA^* SCCs. Genes consistently changing in both, *Hras^G12V^* and *Sox2^KO^*, models are shown. p = exact hypergeometric probability **(D)** Heatmap illustrates z-score transformed FPKM values of genes that are differentially bound and expressed dependent on SOX2-phosphorylation in SCCs. Genes are ranked based on their mean expression level and their up-regulation in SOX2^DD^ or SOX2^AA^ expressing tumors. *Klf4* surfaced as the highest expressed, direct SOX2 target gene whose expression increases significantly in both of our SOX2^DD^ compared to SOX2^AA^ expressing SCC models. **(E-F)** Confocal microscopy reveals significantly increased KLF4 staining in *Sox2^KO^;F-SOX2^DD^* compared to *Sox2^KO^;F-SOX2^AA^* tumor sections **(E)** and box-plots illustrate the differences in KLF4 staining densities. n=3 tumors and 2 fields/tumor. p = unpaired parametric two tailed t-test.

However, differential read count analyses identified a small subset of cis-regulatory elements that were preferentially F-SOX2^AA^ or F-SOX2^DD^ bound (Figure 4A-B). We used the genomic regions enrichment of annotations tool (GREAT, (McLean et al., 2010)) to identify genes with a transcriptional start site within 1 Mb of each SOX2 binding site and intersected them with genes that are consistently up- or down-regulated in *Hras^G12V^;F-SOX2^AA^* and *Sox2^KO^;F-SOX2^AA^* compared to *Hras^G12V^;F-SOX2^DD^* and *Sox2^KO^;F-SOX2^DD^* tumors, respectively. Surprisingly, this approach identified only four genes that were inhibited and four genes that were activated by increased SOX2^AA^ compared to SOX2^DD^ binding (Figure 4C). *Klf4* emerged as the gene with the highest average expression, and it was consistently suppressed in SOX2^AA^ compared to SOX2^DD^ SCCs in both of our tumor models (Figure 4D). Quantitative confocal immuno-fluorescence microscopy confirmed KLF4 expression is significantly increased in basal and suprabasal *Sox2^KO^;F-SOX2^DD^* compared to *Sox2^KO^;F-SOX2^AA^* SCC cells (Figure 4E-F), similar to what we observed in *Hras^G12V^;F-SOX2^DD^* compared to *Hras^G12V^;F-SOX2^AA^* tumors (Figure 2F).

### Phosphorylated SOX2 is evicted from a specific *Klf4* enhancer to permit KLF4 binding and increased *Klf4* expression

Our ChIP-seq and differential expression data suggest SOX2 inhibits *Klf4* transcription, but its expression can be restored when SOX2 becomes phosphorylated in SCC cells. Indeed, our differential SOX2 ChIP-seq analyses identified F-SOX2^AA^ but not F-SOX2^DD^ on a large enhancer cluster (EC944) located 944 kb distal to the *Klf4* transcriptional start site (TSS) (Figure 5A). H3K27Ac ChIP-seq also indicated that this enhancer is more active in *Sox2^KO^; SOX2^DD^* compared to *Sox2^KO^; SOX2^AA^* SCCs and we found Mediator (Med12) enrichment at this site, suggesting promoter-enhancer interactions. To test this hypothesis, we used chromatin conformation capture sequencing (4C-seq) with a bait located at the *Klf4* TSS to unbiasedly search for promoter-enhancer interactions (Figure 5A). 4C-ker (Raviram et al., 2016) identified statistically significant interactions between the *Klf4* promoter and several cis-regulatory elements including *Klf4^EC944^* and *Klf4^EC390^*, which we previously identified as a potential SOX2 and PITX1 bound *Klf4* enhancer (Sastre-Perona et al., 2019). However, *Klf4^EC390^* was, in contrast to *Klf4^EC944^*, equally bound by SOX2^AA^ and SOX2^DD^.

**Figure 5.**
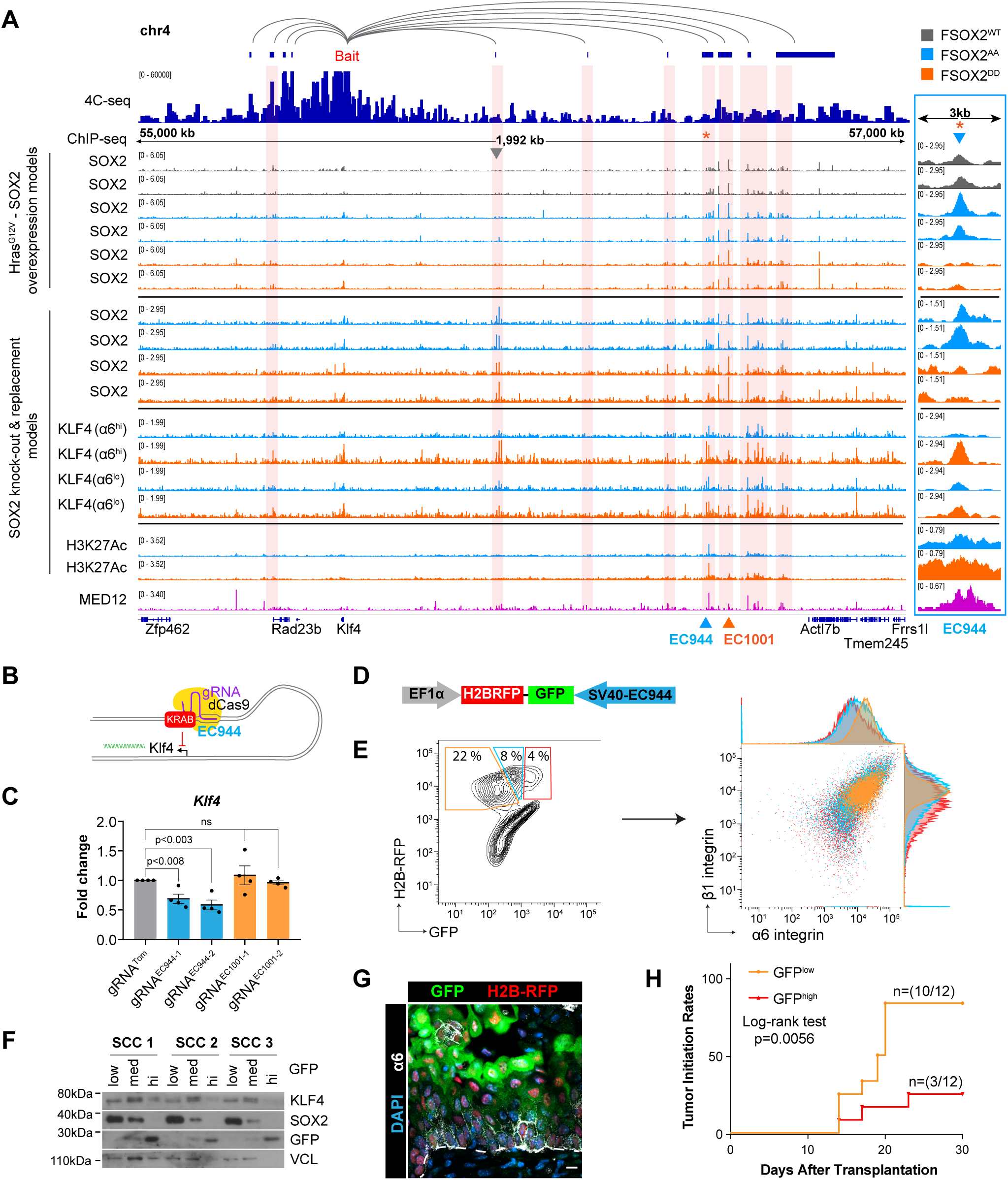
Phosphorylated SOX2 is evicted from a Klf4 enhancer to restore *Klf4* expression. **(A)** 4C-seq and SOX2, KLF4, CTCF, MED12, and K27Ac ChIP-seq tracks identify Klf4 enhancers. 4C-ker visualizes significant interactions between a bait at the Klf4 promoter and cis-regulatory elements by 4C-seq (pink highlights). ChIP-seq signals – significantly (p<0.05) enriched in F-SOX2^AA^ (blue triangle) or F-SOX2^DD^ (orange triangle) – identify putative SOX2-phosphorylation regulated enhancers. Orange asterisk marks site with differential KLF4 enrichment (p<0.05) in ɑ6^hi^/β1^hi^ SCC cells between *SOX2^KO^;F-SOX2^DD^* and *SOX2^KO^;F-SOX2^AA^* SCCs. **(B)** Model illustrating *Klf4* repression upon CRISPRi targeting of *Klf4^EC944^*. **(C)** *Klf4* mRNA fold change after CRISPR-interference (CRISPR-i), targeting either *Klf4^EC944^* or *Klf4^EC100^*. Mean +/- s.e.m. n=4. p = unpaired parametric two-tailed Student’s t-test. **(D)** Schematic of EC944 reporter construct. H2B-RFP is constitutively expressed under an EF1a promoter while GFP expression is controlled by *Klf4^EC944^*. **(E)** Representative contour plot of reporter-containing SCCs separating into GFP low (orange), medium (blue), and high (red) populations **(left)** and scatter plot with adjunct histograms illustrating the inversely proportional expression of *Klf4^EC944^* reporter and self-renewal marker α6-integrin **(right)**. **(F)** Western blotting of protein lysates from sorted reporter^+^ SCCs show SOX2 levels decrease with GFP expression while KLF4 increases. **(G)** Fluorescence microscopy image showing *Klf4^EC944^*-GFP reporter expression. Increased GFP expression is detected in more differentiated suprabasal SCC cells. Scale bars = 10µm. **(H)** Tumor initiation rates from serial transplantation of GFP^Low^/RFP^Hi^ or GFP^Hi^/RFP^Hi^ cells we identified with the *Klf4^EC944^*-GFP reporter. p = Mantel-Cox Log-rank test.

To functionally test if EC944 regulates *Klf4* transcription, we designed two independent *gRNAs* that targeted either a SOX2 or KLF4 motif located at the peak summit **(Figure S4A).** A *gRNA* directed against tdTomato (*gRNA^Tom^*) served as control. Induction of HA-dCas9-KRAB with doxycycline significantly reduced *Klf4* transcription when it was targeted to *Klf4^EC944^* compared to *gRNA^Tom^* or two *gRNAs* targeting SOX2 and KLF4 bound sites at EC1001 (Figure 5B-C). To further characterize *Klf4^EC944^* we cloned it into a fluorescent activity reporter (Figure 5D). Next, we transduced TPCs with this reporter, transplanted them into mouse dermis and assessed the expression of *Klf4^EC944^* driven GFP expression within transduced cells that constitutively expressed H2B-RFP. FACS analyses distinguished cells with low, medium and high GFP expression within the RFP^+^-lineage and their GFP expression levels correlated inversely with ɑ6- and β1-integrin levels (Figure 5E). Furthermore, SOX2 expression declined with increasing GFP expression, whereas KLF4 levels transiently increased (Figure 5F). Consistent with these data we detected only a few basal SCC cells with weak GFP expression, while GFP was highly expressed in supra-basal SCC layers (Figure 5G) and RFP^hi^/GFP^low^ SCC-cells initiated tumors more efficiently than RFP^hi^/GFP^hi^ SCC-cells, when they were transplanted into mouse dermis (Figure 5H). Collectively, these data suggest that SOX2 binds *Klf4^EC944^* to repress *Klf4* transcription in TPCs, but when SOX2 is phosphorylated, it is specifically evicted from this site resulting in increased KLF4 expression and squamous differentiation.

Because *Klf4^EC944^* contains two KLF4 motifs directly adjacent to SOX2 and PITX1 motifs (**Figure S4A**), we wondered whether the eviction of SOX2 upon phosphorylation would allow KLF4 to occupy this enhancer and convert it from a transcriptional repressor into an activator. To test this hypothesis, we isolated ɑ6^hi^/β1^hi^ and ɑ6^lo^/β1^lo^ cells from lineage marked *SOX2^KO^;F-SOX2^AA^* and *SOX2^KO^;F-SOX2^DD^* SCCs by FACS (**Figure S4B**) and determined the occupancy of KLF4 in these cells with ChIP-seq. We found KLF4 is hardly detected on *Klf4^EC944^* in ɑ6^hi^/β1^hi^ *SOX2^KO^;F-SOX2^AA^* SCC cells, but it was significantly enriched at this site in ɑ6^hi^/β1^hi^ *SOX2^KO^;F-SOX2^DD^* SCC cells (Figure 5A). KLF4 was also detected at comparable levels in ɑ6^lo^/β1^lo^ cells of *SOX2^KO^;F-SOX2^AA^* and *SOX2^KO^;F-SOX2^DD^* SCCs. These data suggest that SOX2 and KLF4 bind to *Klf4^EC944^* in a mutually exclusive manner, SOX2 dominates over KLF4 in TPCs, and it inhibits *Klf4* transcription and KLF4 dependent differentiation. However, when SOX2 becomes phosphorylated, it is evicted from *Klf4^EC944^*, which allows KLF4 to bind, activate the enhancer, and boost KLF4 expression and squamous differentiation.

### Increased KLF4 expression and chromatin binding promotes epidermal differentiation

As a result of increased KLF4 expression, we detected significantly more chromatin bound KLF4 in both ɑ6^hi^/β1^hi^ and ɑ6^lo^/β1^lo^ fractions of *SOX2^KO^;F-SOX2^DD^* compared to *SOX2^KO^;F-SOX2^AA^* SCCs with ChIP-seq (Figure 6A). *De novo* motif predictions identified KLF4-like motifs in >60% of KLF4 ChIP-seq peaks at their peak summits (Figure 6A**, S4C**). Although differential KLF4 binding data revealed 290 sites with KLF4 enrichment in ɑ6^hi^/β1^hi^ cells of *SOX2^KO^;SOX2^AA^* compared to *SOX2^KO^;SOX2^DD^* SCCs, its enrichment was rather modest. In contrast, we identified 317 sites that were enriched in ɑ6^hi^/β1^hi^ cells of *SOX2^KO^;SOX2^DD^* compared to *SOX2^KO^;SOX2^AA^* SCCs with an ∼3-fold median enrichment in chromatin binding (Figure 6B). Intersection of these KLF4 bound sites with transcripts that are significantly up-regulated in *Sox2^KO^;SOX2^AA^* or Sox2^KO^;SOX2^DD^ SCC cells (Figure 6C) revealed 138 genes that are directly upregulated as a response of increased KLF4 expression in *Sox2^KO^;SOX2^DD^* SCCs. Amongst these KLF4 targets we found *Klf4* itself along with genes that promote epidermal differentiation, keratinization, and barrier formation (Figure 6D-E). Conversely, we also noticed increased KLF4 enrichment on the SCC specific SOX2 enhancer in SOX2^DD^ mutant SCCs. These data suggest KLF4 drives the expression of squamous differentiation genes (Figure 6E) and it feeds back and occupies the KLF4-motif in the SCC specific *Sox2* enhancer (Figure 6F,G) to inhibit SOX2 expression (Sastre-Perona et al., 2019) and stabilize the differentiated, post-mitotic SCC-cell state.

**Figure 6.**
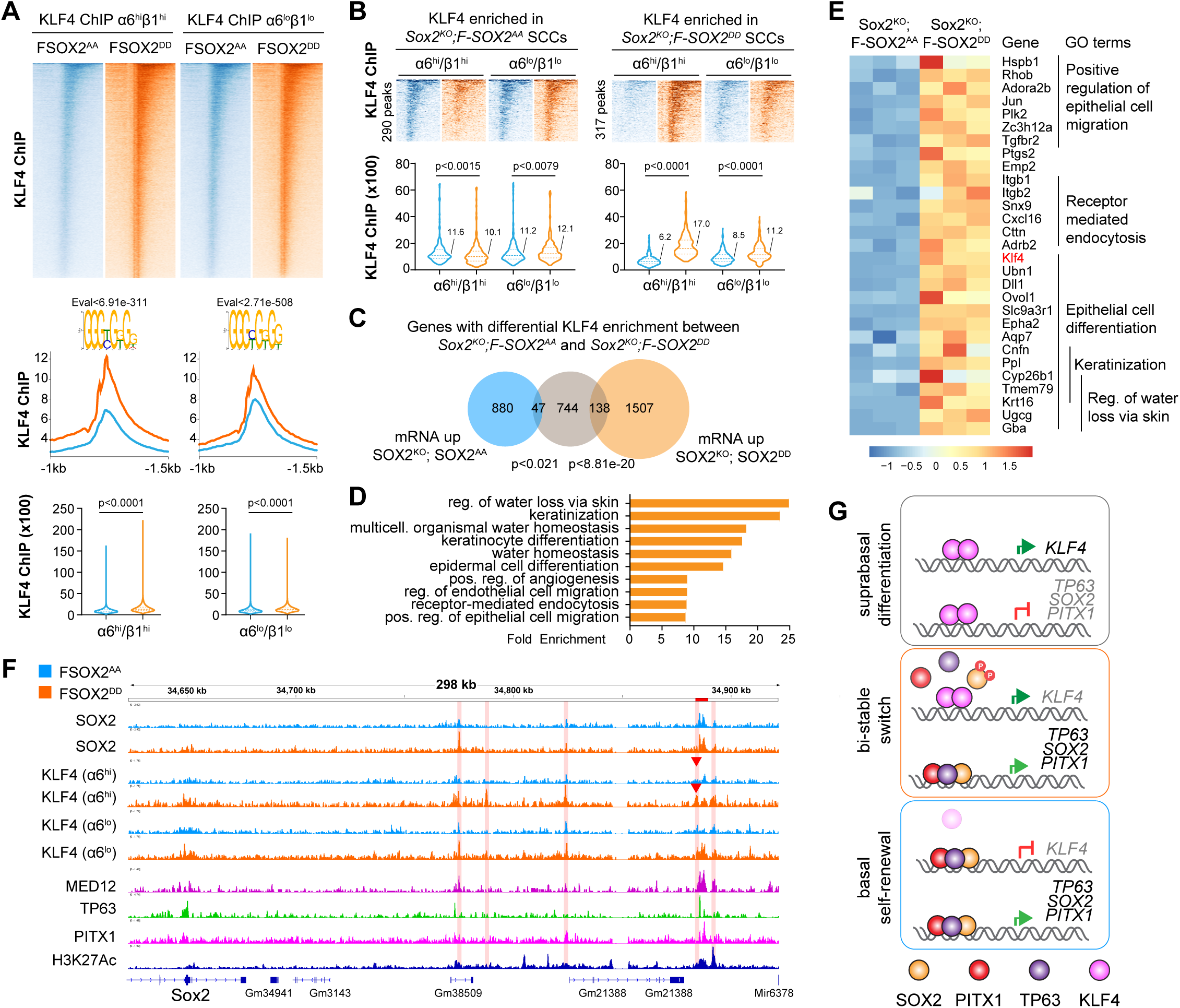
Increased KLF4 expression drives differentiation in F-SOX2^DD^ SCCs. **(A)** Heatmaps, histograms and violin plots of KLF4-like motif containing KLF4 ChIP-seq peaks of FACS-isolated ɑ6^hi^/β1^hi^ and ɑ6^lo^/β1^lo^ SCC cells reveal increased KLF4 occupancy in *Sox2^KO^;F-SOX2^DD^* compared to *Sox2^KO^;F-SOX2^AA^* tumors. DREME identified transcription factor motif at the peak summit with corresponding E-value. p = unpaired non-parametric Mann-Whitney t-test. **(B)** Heatmaps and violin plots (median values listed) of KLF4 ChIP-seq peaks with differential enrichment in ɑ6^hi^/β1^hi^ SCC cells of *Sox2^KO^;F-SOX2^AA^* or *Sox2^KO^;F-SOX2^DD^* tumors. n = 290 peaks (SOX2^AA^) and n = 317 peaks (SOX2^DD^). p = Mann-Whitney t-test. **(C)** Intersection of genes associated with KLF4 bound sites that were significantly enriched in *SOX2^KO^;F-SOX2^AA^* or *Sox2^KO^;F-SOX2^DD^* SCCs with differentially expressed transcripts from these tumors. p = exact hypergeometric probability. **(D)** GO analysis of the genes with differential KLF4 ChIP-seq enrichment and increased expression in *Sox2^KO^;F-SOX2^DD^* tumors. **(E)** Heat map representation of differentially expressed transcripts and their corresponding GO categories. **(F)** ChIP-seq tracks show increased KLF4 occupancy at the KLF4-motif (red arrowhead) containing *Sox2* enhancer site that inhibits *Sox2* expression. **(G)** Working model: SOX2 inhibits *Klf4* to maintain TPC self-renewal in basal SCC cells. SOX2 phosphorylation evicts SOX2 from a bi-stable *Klf4* enhancer switch allowing residual KLF4 to bind and boost its own expression. Increased KLF4 levels promote squamous differentiation.

## Discussion

The fate choice between stem cell self-renewal and differentiation is regulated by bistable transcriptional networks that allow cells to stably reside in one of two alternative cell states (Davis and Rebay, 2017; Young, 2011). Genetic gain- and loss-of-function studies identified master regulatory transcription factors as the core components of these networks and they found these transcription factors promote self-renewal by enhancing each other’s expression in positive feed-forward circuits, while they repress transcriptional programs that promote differentiation. Conversely, differentiation promoting transcription factors enhance their own expression as they inhibit the self-renewal network. We and others previously identified several components of a bistable transcriptional network that governs self-renewal and differentiation in SCCs (Boumahdi et al., 2014; Sastre-Perona et al., 2019; Siegle et al., 2014). We found this network is comprised of mutually exclusive SOX2-PITX1-ΔNp63 self-renewal and KLF4 dependent differentiation circuits (Sastre-Perona et al., 2019). Likewise, a cancer driving SOX2-ΔNp63-KLF5 feed-forward circuit has been described in human esophageal SCCs (Jiang et al., 2018, 2020; Watanabe et al., 2014). Although new components of this SCC regulatory bistable transcriptional network are being identified, it remained unclear how TPCs can differentiate in steady state tumors, if KLF4 controls the switch from self-renewal to differentiation, but its expression is blocked in self-renewing TPCs.

In this study we identified a molecular mechanism that resolves this self-renewal – differentiation paradox. We discover that SOX2 – a core component of the self-renewal circuit – can get phosphorylated in TPCs and increased SOX2 phosphorylation reduces TPC self-renewal and it enables them to differentiate into post-mitotic SCC cells (Figure 6G). Although we found that changes in SOX2 phosphorylation affected TPC self-renewal and differentiation rates and that these changes had a profound impact on SCC initiation and tumor growth, we found no changes in cell proliferation and survival programs that have been linked to SOX2 in SCCs (Boumahdi et al., 2014). This finding is consistent with a prior study that found cyclin dependent kinase mediated SOX2 phosphorylation of S37 and S251 enhances the ability of SOX2 to induce a pluripotent embryonic stem cell (ESC) state, and that SOX2 phosphorylation was dispensable for ESC self-renewal (Ouyang et al., 2015). These studies suggest phosphorylated SOX2 has highly specific and context dependent functions, and a better understanding of their roles and regulation could inform the development of differentiation therapies for patients with SCCs or other SOX2 regulated cancers like glioblastoma or astrocytoma (Modrek et al., 2017; Suvà et al., 2014).

Although several post-translational SOX2 modifications have been reported, their roles and regulation are context specific and poorly understood (Williams et al., 2019). While human SOX2 S37 and S251 phosphorylation could have simply affected nuclear import or export, reduced SOX2 stability, or impaired its DNA binding ability, we found its effects are much more context dependent. Quantitative ChIP-seq studies with non-phosphorylated SOX2^AA^ and phosphorylation mimetic SOX2^DD^ mutants with spike in controls for data normalization revealed that their DNA binding patterns are highly concordant, they only differed in ∼5% of SOX2^AA^ or SOX2^DD^ bound sites, where SOX2 phosphorylation increased or decreased SOX2 – DNA interactions dependent on the gene regulatory site. We only identified eight gene regulatory sites that were more strongly bound by SOX2^AA^ compared to SOX2^DD^ and that could be linked to consistent changes in gene expression in both of our tumor models. The expression of half of these genes increased in SOX2^DD^ tumors, while the other genes decreased. No consistent changes in gene expression were observed in genes that were more strongly bound by SOX2^DD^.

Amongst the gene regulatory elements that were bound by SOX2^WT^ and SOX2^AA^ but not SOX2^DD^ we identified an enhancer cluster *Klf4^EC944^* that is located 944 kb upstream the *Klf4* TSS. Although this enhancer is much closer to *Actl7b* than to *Klf4* in linear distance, we were unable to detect *Actl7b* in SOX2^AA^ or SOX2^DD^ expressing SCCs, while the expression of *Klf4* was significantly higher in SOX2^DD^ compared to SOX2^AA^ SCCs. Consistent with these changes in gene expression and SOX2 binding, we found the *Klf4* promoter interacts with EC944 by 4C-seq. Additionally, the activity of a LV-PGK-H2B-RFP;EC944-GFP expression reporter increases as ɑ6/β1-integrin and SOX2 levels decline, and KLF4 expression strengthens while cells transition from basal- to suprabasal SCC layers. These data suggest SOX2 binds to EC944 to inhibit *Klf4* transcription in TPCs, but SOX2 becomes evicted from this site when it is phosphorylated and this allows residual KLF4 to bind this site to enhance its own expression. Once KLF4 expression increases it begins to accumulate at many gene regulatory elements across the epigenome to promote the expression of squamous differentiation markers and inhibit TPC self-renewal and proliferation.

Although this study uncovers a novel gene regulatory mechanism that can resolve the self-renewal differentiation paradox, it also reveals that *de novo* expressed SOX2 promotes clonal expansion and SCC growth partly because it can directly inhibit the expression of several TGFβ- and Notch-pathway components with known tumor suppressive functions in stratified epithelial cells. However, our study also raises new questions. One of the most exciting emerging questions is why phosphorylated SOX2 gets evicted from *Klf4^EC944^*, but not from most other SOX2 bound sites? It is intriguing to speculate that the addition of negatively charged phosphate groups could introduce structural changes that perturb the interaction of SOX2 with other transcription factors or transcriptional co-regulators. These changes could weaken their protein-DNA interactions and it may allow residual KLF4 to displace SOX2 from sites that are rich in KLF4 binding motifs. Indeed, *Klf4^EC944^* contains central PITX- and SOX-motifs that are surrounded by several KLF-motifs and it is possible that SOX2 and KLF4 competitively occupy this site, where their binding kinetics would depend on the active concentrations of SOX2, KLF4, and their respective transcriptional partners.

Another emerging question is which kinases and phosphatases are responsible for the phosphorylation of SOX2 in SCCs. In vitro studies suggested that S37 and S251 of human SOX2 can be phosphorylated by CDK2, Aurora kinase A, or MAPK1/3 (Ji et al., 2018; Lim et al., 2017; Ouyang et al., 2015), but it remains to be tested whether SOX2 is a substrate of cell cycle regulated or rather transcriptional CDKs in SCCs. Even more exciting than the identification of the SOX2 regulatory kinases would be the identification of SOX2 regulatory phosphatases as their inhibition would increase SOX2 phosphorylation and squamous differentiation in tumors thus inhibiting SCC growth. Although these questions remain to be answered in future studies, our data suggest that SOX2 may cycle between phosphorylated and un-phosphorylated states and this cycling behavior along with the competitive occupancy of *Klf4^EC944^* by SOX2 or KLF4 may explain why the fate choice between self-renewal and differentiation is stochastically regulated (Driessens et al., 2012) and how cell fate decissions can become biased towards self-renewal or differentiation dependent on the effective concentration of SOX2 and KLF4, respectively.

## Acknowledgments

We thank E. Fuchs and P. Khavari for reagents, Ramya Raviram and Sana Badri for computational advice, and Eva Gonzalez, Eva Hernando-Monge and Timothee Lionnet for discussions. We are grateful to the NYU Langone Health Flow cytometry core facility, the Division of Comparative Medicine for expert handling and care of mice, the NYU Langone Health genome technology center for next generation sequencing, and the High-Performance Computing Facility for cluster access and data storage. This research was supported by NIH grant 1R01-CA248175-01A1, American Cancer Society grant RSG-16-033-01-DDC, Worldwide Cancer and the Orbuch-Brand Pilot Grant Program for Cancers of the Skin to M.S., T32 CA009161 and T32 AR064184 to A.S-P. and T32 CA009161 to S.H-P. Core funding was partially supported by NIH grant P30 CA016087 to the Perlmutter Cancer Center.

## Author Contributions

Conceptualization, S.H-P., A.S-P. and M.S.; Methodology, S.H-P., M.A., A.S-P., P.R., Z.Y., B.A-O., and M.S.; Formal Analysis, S.H-P., M.A., A.S.-P., Z.Y. and M.S.; Investigation, A.S.-P., S.H-P., M.A., J.S., I.A., S.B., and M.S.; Writing—Review and Editing, S.H.P. and M.S.; Writing— Review and Editing, S.H-P., A.S-P., Y.Z., S.B. and M.S.; Visualization, S.H-P., A.S-P. and M.S.; Funding Acquisition, M.S.

## Declaration of Interests

The authors declare no competing interest.

## Supplementary Figures

**Figure S1.**

**(A)** Work flow of experimental approach to identify F-SOX2 phosphorylation sites with tandem mass spectrometry (MS/MS).

**(B)** Silver stained gel showing 5% of F-SOX2 immuno-precipitate used for MS/MS.

**(C)** Base peak chromatogram for F-SOX2

**(D-E)** Spectral traces of peptides identifying phosphorylated Serine-37 **(D)** and Serine-251 **(E)**.

**(F)** Confocal micrograph of patient SCC stained with pSer^251^-SOX2 and SOX2.

**(G)** Phos-TAG western blot of human SCC protein lysates transduced with *F-SOX2^WT^* or *F-SOX2^AA^* transgenes showing F-SOX2^AA^ is, in contrast to F-SOX2^WT^, not phosphorylated. Black arrows mark F-SOX2 bands. Grey arrows mark non-specific bands that are also detected in un-transduced cells.

**(H)** FACS-gating strategy to determine GFP:RFP ratio in clonal competition experiments.

**Figure S2.**

**(A)** FACS-gating strategy to isolate epidermal progenitor cells (Sca1+) and hair follicle stem cells (CD34+) from mouse back skins.

**(B)** Western blot of epidermal progenitor cells transduced with *Hras^G12V^*, *Hras^G12V^;F-SOX2^WT^, Hras^G12V^;F-SOX2^AA^*, or *Hras^G12V^;F-SOX2^DD^* showing transgene but no endogenous SOX2 expression.

**(C)** FACS-gating strategy to sort out lineage positive cells (green, GFP+ gate) for RNA-sequencing. Gating strategy for determining relative abundance of ɑ6^hi^/β1^hi^ and ɑ6^lo^/β1^lo^ fractions are also shown. Gates are defined based on ɑ6- and β1-integrin expression values of the GFP-negative, normal skin epithelial cell lineage, which serves as an internal control to define top and bottom 25 % gate limits.

**(D)** *Sox2* FPKM values in *Hras^G12V^*, *Hras^G12V^;F-SOX2^WT^, Hras^G12V^;F-SOX2^AA^*, or *Hras^G12V^;F-SOX2^DD^* SCCs. Red dot denotes *Hras^G12V^* tumor with de novo expression of endogenous Sox2. Mean +/- s.e.m. n=3. Note: FPKM values for *Hras^G12V^;F-SOX2^WT^, Hras^G12V^;F-SOX2^AA^*, or *Hras^G12V^;F-SOX2^DD^* tumors are for the transgene and not endogenous *Sox2*.

**(E)** Confocal micrographs of tumor sections stained with pS251-SOX2 and SOX2 antibodies.

**(F)** Schematic of BrdU/EdU double label incorporation self-renewal assay. Mice are injected with EdU (0h) and BrdU (2h) before sacrificing at 8h after initial EdU injection. >95% of EdU+ only cells complete one round of cell division in the 6h between BrdU injection and sacrifice. Tracking EdU+ only cells (differentiated, K10+ or self-renewing, K10-) allows calculation of self-renewal rates.

**Figure S3.**

**(A)** Bar graphs showing percentages of recovered *SOX2^KO^;F-SOX2^WT^, SOX2^KO^;F-SOX2^AA^, and SOX2^KO^;F-SOX2^DD^* clones.

**(B)** FACS-gating strategy to determine relative α6-integrin expression. Gates are kept constant for all samples.

**Figure S4.**

**(A)** CRISPRi *gRNA* targeting schematic. Independent *gRNA’s* were designed to target either SOX2 or KLF4 motifs within the ChIP-seq peak for the *F-SOX2^AA^* (top) or *F-SOX2^DD^* (bottom) bound peaks in EC944 and EC1001, respectively.

**(B)** FACS-gating strategy to isolate ɑ6^hi^/β1^hi^ and ɑ6^lo^/β1^lo^ SCC cells from *SOX2^KO^;F-SOX2^AA^, and SOX2^KO^;F-SOX2^DD^* tumors for KLF4 ChIP-seq experiments.

**(C)** Stacked bar graphs illustrate percentage of peaks with KLF4 like transcription factor motifs we discovered de-novo with DREME and their distribution in respect to the closest transcriptional start site.

## Materials and methods

### Resource Availability

Further information and requests for resources and reagents should be directed to and will be fulfilled by the Lead Contact, Markus Schober.

### Materials Availability

Plasmids generated in this study are available upon request.

### Data and Software Availability

This study did not generate any new code. Data sets have been uploaded to NCBI GEO with the accession number GSE158207.

### Experimental Model and Subject Details

#### Cell lines

Primary murine SCC-TPC lines were established from malignant tumors induced with DMBA following the previously described complete carcinogenesis protocol (Guasch et al., 2007; Schober and Fuchs, 2011; Siegle et al., 2014). Human A431 cells were grown in DMEM supplemented with 10% fetal bovine serum (FBS). Human SCC25 cells were grown in P-media (DMEM-F12 3:1 media (US Biologicals), sodium bicarbonate (Sigma), L-glutamine (Invitrogen) and Pen/Strep solution (Invitrogen)) supplemented with 10% FBS. Differentiation was induced in primary cells by increasing [Ca^+2^] from 0.05 mM to 1.5 mM for the times indicated.

#### Mice

6 week old female nude (NU/NU [088] Charles River) mice were used for all orthotopic allo/xenograft studies. *Hras^G12V/G12V^* (Chen et al., 2009) and *Rosa26^YFP/YFP^* Cre-reporter mice (Jackson Laboratories) on C57Bl/6 background were used for in utero transduction experiments and equal numbers of male and female mice have been analyzed in tumor formation studies. Tumors were detected by palpation, measured with digital calipers (in mm), and volume calculated with the equation: 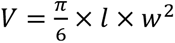. All animal experiments were performed in accordance with the guidelines and approval by the Institutional Animal Care and Use Committee at New York University Langone Health.

## Method Details

### Constructs

Human SOX2 was PCR amplified from genomic DNA we isolated from human A431 cells using primers with the first half of the p2A tag sequence in the 3’ primer. eGFP was amplified from LZRS-IRES-eGFP (Addgene 21961) using primers with the second half of the p2A tag sequence in the 5’ primer. Both PCR products were purified and used as the template for a second round of PCR amplification to generate the SOX2-p2A-eGFP sequence, which was first cloned into the Zero Blunt TOPO cloning vector (ThermoFisher 450245), and then subcloned into a pcDNA 3.1 vector containing the flag-flag-streptavidin-streptavidin (FFSS) tag (generous gift from Dr. Michele Pagano) using the EcoRV restriction site. The full FFSS-SOX2-p2A-eGFP (FSOX2) sequence was excised from the pcDNA3.1 vector with the restriction enzymes NheI and EcoRV and ligated into pLKO.1 (Addgene 10878). FSOX2^AA^ and FSOX2^DD^ mutant constructs were obtained by sequential site directed mutagenesis and sequenced to ensure no other mutations were present.

The p2A-H2B-GFP sequence was amplified from pLKO Histone H2B-GFP (Addgene 25999) and the p2A-H2B-RFP sequence was amplified from pLKO Histone H2B-mRFP1 (Addgene 26001) with primers containing the p2A sequence at the 5’ primer. Subsequent steps to generate FFSS-SOX2-p2A-H2B-eGFP or - FFSS-SOX2-p2A-H2B-mRFP constructs were the same as described above.

For in utero transduction experiments, human SOX2 from the CCSB Broad ORF library (Yang et al., 2011) was cloned into the pLEX_307 (Addgene 41392) vector. SOX2^AA^ or SOX2^DD^ constructs were generated by site directed mutagenesis with PCR using the human SOX2^WT^ plasmid as template.

pLKO-HRas^G12V^-Puro was generated by subcloning HRas^G12V^ from PQCXIX-PGK-Hras^V12^- IRES-H2B-RFP (generous gift from Dr. Paul Khavari) into the pLKO vector.

To make the guide RNA backbone, the puromycin resistance gene was PCR amplified from the pLKO.1 vector and cloned into the pEHL958-pLKO-GFP-sgRNA backbone (generous gift from Dr. Eva Hernando-Monge), by excising GFP with BamHI and SalI and replacing it with the puromycin resistance gene. Individual guide RNAs were designed with the Benchling CRISPR gRNA tool and cloned into this vector using the BbsI restriction site.

The EC944-eGFP reporter was constructed by PCR amplification of mouse genomic DNA with a primer pair that introduced KpnI and BsaBI restrictions sites and it was cloned into corresponding sites of a lentiviral Notch-reporter (kind gift from Scott Williams (Williams et al., 2011)).

### Cell culture

Primary SCC cells were infected with VSV-G psuedotyped lentiviruses. Lentiviruses were produced by transfecting 293FT cells with pLKO sgRNA or FSOX2 expression vectors and helper plasmids pMD2-VSVg and pPAX2 (Addgene plasmid 12259 and 12260) using Lipofectamine 2000 (Thermo Fisher). 293FT cells were cultured in DMEM (Gibco) media supplemented with 10% FBS and Pen/Strep solution. Virus containing media was harvested 48h and 72h post-transfection and filtered with 0.45µm pore size mixed cellulose esters syringe filters (EMD Millipore SLHA033SS). For lentiviral infection, 3 × 10^4^ cells per well were plated into a 6 well plate, incubated with a 1:3 dilution of viral supernatant with 30ug/mL of polybrene and spun at 1,100 × g for 30 min at 37°C. For FSOX2 stable cell lines, eGFP+ cells were selected by FACS using a BD FACSAria II equipped with 405, 488, 561, 647, and 355nm lasers. Primary epidermal progenitor cells (CD49f+/Sca1+/ CD34-) were extracted from 21 day old mice with trypsin, isolated by FACS and cultures were established on 3T3 feeder layers. Cells were transduced with pLKO-HRas^G12V^-Puro on 3T3 feeder layers. Transduced cells were selected with 1µg/mL puromycin (Sigma-Aldrich) containing media for 2 days before they were transduced with pLKO-FSOX2-p2A-eGFP, pLKO-FSOX2^AA^-p2A-eGFP, or pLKO-FSOX2^DD^-p2A-eGFP and sorted for eGFP expression.

### CRISPR/Cas9 knockouts

For SOX2^KO^ –SOX2 replacement experiments, primary SCC cultures were first transduced with pLKO-FSOX2^AA^-p2A-eGFP or pLKO-FSOX2^DD^-p2A-eGFP before the entire *Sox2* coding region was deleted using two gRNAs targeting the 5’ and 3’ UTR’s. Guide RNA’s were chosen based on similar cutting efficiency scores and to minimize off targets using the Benchling CRISPR gRNA design tool. Cells were infected with sgRNAs, selected with 1ug/mL puromycin (Sigma-Aldrich) containing media for 2 days before they were infected with lenti-Cas9-Blast (Addgene 52962) and selected with 5ug/mL blasticidin for 3 days. After selection, 100 cells were plated in a 10cm dish and single clones were picked with 6mm cloning cylinders when clone sizes reached ∼100 cells. Clones were screened by PCR amplification of the Sox2 locus with primers flanking the Sox2 coding sequence using gDNA as template as well as western blotting, where endogenous SOX2 and FSOX2 can be distinguished based on the FFSS-tag size difference.

### CRISPRi

Independent gRNAs targeting either the SOX2 or KLF motif within peaks EC944 or EC1001 were designed using the Benchling gRNA design tool (www.benchling.com). TPC cell lines were infected with rTTa-N144 (Addgene #66810) and selected with hygromycin (10ug/mL). The cells were then infected with TRE-dCas9-KRAB-IRES-GFP (Addgene #85556), induced with doxycycline (1µg/mL) for 48h, and FACS sorted for GFP+ cells. The GFP^+^ population was expanded and lentivirally infected with the gRNAs before selection with puromycin (2ug/mL). dCas9-KRAB expression was induced with doxycycline (1µL/mL) for 48h before harvesting cells for protein and RNA to check for changes in *Klf4* expression. KRAB-dCas9-HA expression was confirmed by western blotting using anti-HA antibodies.

### Clonal competition

Lentiviruses containing FSOX2-p2A-H2B-eGFP or FSOX2-p2A-H2B-mRFP were generated by mixing equimolar concentrations of both plasmid DNAs during transfection of 293FTs. Human A431 or primary murine TPCs were infected at low MOI. The percentage of GFP:RFP containing cells were checked on a BD LSRII UV flow cytometer equipped with 405, 488, 561, 647, and 355nm lasers 72h after infection. 1 × 10^5^ cells from pools that contained ∼5% GFP+:RFP+ cells were used for each transplantation.

### Tumor isolation and flow cytometry

Tumors from murine allografts were isolated as previously described (Sastre-Perona et al., 2019). Briefly, tumor tissue was separated from normal skin, blood vessels and connective tissue as much as possible before mincing and incubation in HBSS (Gibco) with 1.25% Collagenase Type IA (Sigma) for 40min at 37°C and shaking in an orbital shaker at 60rpm. 62.5U/mL DNAseI (Worthington) is added afterwards and incubated for another 10min, shaking. The tumor suspensions are filtered through 70µm then 40µm mesh strainers (Thermo Fisher). Remaining undigested tumor pieces are incubated with 0.25% trypsin (Gibco) for 10min, 60rpm before filtering through the mesh strainers again. Cell suspensions are spun down at 300 × g for 10 min at 4°C. The supernatant is discarded and the tumor pellet is washed with 1x RBC lysis buffer (Biolegend), pelleted at 300 × g for 3min, and washed with staining buffer (2% chelexed FBS and 0.5U/mL DNAseI in dPBS).

For flow cytometry, tumor suspensions were stained in the dark for 30min on ice and washed after staining to remove excess antibodies. DAPI (40,6-diamidino-2-phenylindole; D1306, Invitrogen) was used for live/dead cell exclusion. FACS data was recorded on either the BD FACSAria II if sorting was required or the BD LSRIIUV if no sorting was required. Flow cytometry data was analyzed using FlowJo software.

### Serial transplantations

For limited dilution serial transplantation studies, parental tumors were processed as described above and sorted on a BD FACSAria II flow cytometer. Total GFP lineage positive cells were sorted out, serially diluted to obtain 1,000 or 100 cells, mixed with 50% Matrigel solutions, and intradermally injected into nude recipient mice (Sastre-Perona et al., 2019).

### Immunoprecipitation and tandem mass spectrometry

#### Immunoprecipitation (IP)

Human SCC25 cells expressing FSOX2^WT^ were lysed in IP buffer (20mM HEPES pH8.0, 10mM KCl, 0.15mM EDTA, 0.15mM EGTA, 100mM NaCl, 1% NP-40), sheared with an insulin syringe to break the nuclei, and incubated for 30 minutes at 4°C while rotating. Lysates are pelleted at max speed in a table top centrifuge (∼21000rpm) for 10 min, 4°, and the supernatant is combined. Lysate pre-clearing was done with 50µL of IP buffer washed protein A agarose beads (Cell Signaling) for 1h at 4°C, rotating. Protein A beads were pelleted at 4000 rcf for 4 min at 4°C and the supernatant was collected. 50µL Anti-Flag M2 beads (Sigma) were washed twice with IP buffer, added to the supernatant, and incubated overnight while rotating at 4°C. Flag beads were washed 3x with IP buffer, 1x with low salt, 1x with high salt, and 2x with LiCl salt wash buffers (20mM Tris-HCl pH8.0, 2mM EDTA, 0.1% SDS, 1% Triton X100 and 167mM NaCl or 500mM NaCl (low/high salt washes), and 10mM Tris-HCl pH8.0, 1mM EDTA, 1% NP-40, 1% deoxycholic acid, and 250mM LiCl for LiCl salt wash), changing to a new tube every wash to reduce background. 5% of the Flag beads were used to check the concentration and IP efficiency by silver stain before using the rest for mass spectrometry.

#### Mass spectrometry

IP’ed FSOX2^WT^ on Flag beads was reduced with DTT at 57°C for 1h, alkylated with iodoacetic acid at RT in the dark for 45min, and run on NuPAGE 4-12% bis-tris gels (Life Technologies). The gel was stained with GelCode Blue (Thermo Fisher) and the FSOX2 band was excised based on size. Excised gel pieces were destained in a 1:1 v/v solution of methanol and 100mM ammonium bicarbonate, dried, and digested with CNBr in 70% TFA overnight in the dark at RT. Digestion was quenched with 1mL of LCMS water and dried in a SpeedVac before being further digested by 250ng of sequencing grade modified trypsin (Promega). 300µL of 100mM ammonium bicarbonate and 200µL of 1M ammonium bicarbonate was added to the gel pieces and digested overnight. 500µL of R2 20µm Poros beads (Life technologies) in 5% formic acid and 0.2% TFA was added to the sample and shaken for 4h at 4°C. The beads were loaded onto equilibrated C18 ziptips (EMD Millipore) using a microcentrifuge for 30sec at 6000rpm. Gel pieces were rinsed 3x with 0.1% TFA and also loaded onto ziptips. Extracted poros beads were washed with 0.5% acetic acid and peptides eluted by the addition of 40% acetonitrile (ACN) in 0.5% acetic acid followed by 80% CAN in 0.5% acetic acid. The organic solvent was removed using a SpeedVac and the protein reconstituted in 0.5% acetic acid. 1/3 of the sample was analyzed through MS. Peptides were separated using an online LCMS using the autosampler of an EASY-nLC 1000 (Thermo Scientific) and gradient eluted from the column directly to the Orbitrap Elite MS (Thermo Scientific) using a 1hr gradient. High resolution full MS spectra were acquired with a resolution of 60,000, an AGC target of 1 × 10^6^, with a maximum ion time of 200ms, and a scan range of 300-1500m/z. Following each full MS, fifteen data-dependent high resolution HCD MS/MS spectra were acquired. All MS/MS spectra were collected using the following instrument parameters: resolution of 15,000, AGC target of 5 × 10^4^, max ion time of 100ms, one microscan, 2m/z isolation window, fixed first mass of 150m/z, and NCE of 30. MS/MS spectra were searched against the SOX2 sequence using Byonic for phosphorylation modifications.

### In utero injections

Lentiviral transduction *in vivo* via ultrasound-guided *in utero* microinjection was performed as previously reported (Beronja et al., 2010; Ying et al., 2018). Briefly, developmental stage of mouse embryos was determined by Vevo 1100 animal ultrasound imager using MS550D transducer (Visualsonics). 1μl of virus diluted to achieve desired transduction level was injected into amniotic cavity of E9.5 day embryos using a glass capillary needle fitted on Celltram Vario micro injector (Eppendorf).

### Self-renewal assay

We used the previously described assay (Ying et al., 2018) as follows: We first administered EdU to animals, followed by BrdU injection 2 hours later. Six hours later we harvested epidermis and detected EdU, BrdU signal as well as differentiation marker K10 by immuno-fluorescence microscopy. We defined the number of EdU+ only cells as E. We calculated the rate of progenitor cell renewal of the EdU+ only cells (E) using the following equation: rate of renewal = (number of K10- E cells) / (total number of E cells).

### Western blotting

Protein lysates were prepared in RIPA buffer (150mM sodium chloride, 0.1% Triton-X 100, 0.5% SDS and 50mM Tris pH 8 in ddH2O) supplemented with Mini EDTA-free protease inhibitor tablets (04693159001, Roche). Protein concentrations were determined with the Pierce BCA Protein Assay Kit (23225, Pierce). Lysates were boiled with 5x Laemmli buffer (6% SDS, 10% β-mercaptoethanol, 30% glycerol, 0.006% bromophenol blue, 0.188M Tris–HCl) for 10 min at 95°C. Novex Sharp Pre-stained Protein Standard (LC5800 Thermo Fisher) was used for molecular weight markers. 30µg of protein was loaded per lane. Gel electrophoresis was performed using a 10% bis-tris gel run at 120V for 2.5h. Gels were transferred for 1h at 4°C at 100V to a 0.45µm nitrocellulose membrane (GE Healthcare Amersham). Transfer efficiency was determined with Ponceau S staining (0.1% (w/v) Ponceau S in 5% (v/v) acetic acid) and membranes blocked with blocking buffer (5% non-fat dry milk in TBST). Membranes were rinsed with TBST before incubating with primary antibodies in blocking buffer overnight. Membranes were washed 3x with TBST, incubated with HRP-conjugated secondary antibodies in blocking buffer for 1h at RT and washed 3x with TBST. Membranes were developed using Supersignal West Pico or Femto Chemiluminscent substrate (Life Technologies, #34080 add #30095) and exposed to X-ray film (F-9024-8_10, GeneMate) using a Kodak X-Omat 2000A Processor.

Antibodies used for western blotting were SOX2 (1:1,000; Abcam, ab92494), pSOX2 (1:1000; Cell Signaling, 92186), FLAG (1:2000, Sigma, F3165), KLF4 (1:1,000; R&D, AF3158), GFP (1:2000, Abcam, Ab290), Vinculin (1:10000; Sigma, V9131), HRP donkey anti-rabbit IgG (H+L) (1:3000; Jackson, 711-035-152), HRP donkey anti-mouse IgG (H+L) (1:3000; Jackson, 711-035-151), and HRP donkey anti-goat IgG (H+L) (1:3000; Jackson, 705-035-147).

### Immunofluorescence, histology, and imaging

Unfixed tumors were embedded in OCT (Sakura Finetek #4483) and frozen. Sections were cut on a Leica cryostat to a thickness of 10µm and mounted on SuperFrost Plus (Thermo Fisher) slides. Slides were air-dried for 10 min, fixed for 10min with 4% formaldehyde, rinsed with PBS, permeabilized and incubated in blocking buffer (5% normal donkey serum, 1% BSA, 0.3% Triton X-100 in PBS) for 1h at RT, and stained with primary antibody diluted in blocking buffer at 4°C overnight in a humidity chamber. Slides were washed 3x with PBS the next day, incubated with fluorophore conjugated secondary antibodies and Hoechst 33342 (83218, Anaspec) for 1h at RT, washed 3x with PBS, and mounted with ProLong Gold (Thermo Fisher) reagent. Imaging was performed on a Nikon Eclipse TiE microscope or a Zeiss LSM780 confocal microscope. Images were analyzed in ImageJ (Fiji).

For H&E stains, tissue slides were fixed as above, washed 2x with 1x PBS, 1x with H2O, stained in hematoxylin (Thermo Fisher #7231) for 3min, rinsed with H_2_O, and counter-stained with eosin (Thermo Fisher #71204) for between 5-10min. Slides were then rinsed with H_2_O, followed by consecutive washes for 1min each of 90%, 100%, and 100% EtOH before dehydration 2x with xylenes at 3min each and mounting.

Primary antibodies used in immunofluorescence (IF): CD49f (1:200; Biolegend, 313618), SOX2 (1:1000; Abcam, Ab92494), SOX2 (1:500, R&D, AF2018), pSOX2 (1:500; Cell Signaling Technologies, 92186), KLF4 (1:1000; R&D, AF3158), Filaggrin (1:1000; BioLegend, 905801), BrdU MoBU-1 (1:100; Thermofisher, B35128) and K10 (1:1000; Biolegend, Poly19054). Secondary antibodies used in IF: AlexaFluor 488 Donkey α-goat IgG (Thermo Fisher, A11055), AlexaFluor 488 Affinipure Donkey α-rabbit IgG (Jackson, 711-545-152), Rhodamine Red Affinipure Donkey α-rabbit IgG (Jackson, 711-295-152), AlexaFluor 568 Donkey α-goat (Thermo Fisher, A11057).

### Chromatin immunoprecipitation and sequencing

For SOX2 ChIP-sequencing, 1 × 10^7^ cells were fixed for 10 minutes with 1% formaldehyde in PBS while rotating at RT and quenched with 0.125M glycine for another 5 min while rotating at RT. After washing twice with PBS, cells were collected and chromatin prepared according to the ChIP-IT High Sensitivity (Active Motif, 53040) protocol. Briefly, cell pellets were re-suspended in 10mL of chromatin preparation buffer supplemented with protease inhibitor cocktail (PIC) and PMSF, nuclei were broken with a dounce homegenizer on ice, and chromatin sheared with a Diagenode Bioruptor 300 at maximum intensity for 20x cycles of 30sec on and 30sec off. A small aliquot of sonicated chromatin was used to confirm sheared chromatin was between 200-1200bp in size, according to manufacturer protocol. IP was performed overnight at 4°C on 30µg of sheared chromatin using 4µg of either SOX2 (R&D, AF2018) or H3K27Ac (Abcam, ab4729). 3µg (10%) human SCC chromatin was included in the IP as an independent spike-in normalization control. Immunoprecipitated samples were washed, de-crosslinked, and DNA purified with a spin column and eluted in 37µL of DNA Purification Elution buffer.

Libraries were prepared by concentrating eluted DNA in a SpeedVac and reconstituting in 12µL of Molecular Biology Grade Water (Corning, 46-000-Cl). 2µL of the solution were used to determine the DNA concentration on a Qubit 2.0 fluorometer (Thermo Fisher). Libraries were prepared using the SMARTer ThruPLEX DNA-Seq kit (Takara Bio, R400675) using barcode indexes from the SMARTer DNA Single Index Kits (Set A or Set B, R400695 or R400697). Library concentration and quality was assessed by Tapestation after the first round of PCR amplification and amplified for 2-3x more cycles if necessary, before AMPure XP bead (Beckman Coulter, A63880) cleanup to remove excess barcodes. Library concentration and quality was assessed a second time by Tapestation before equimolar amounts of each sample were multiplexed and run on either an Illumina HiSeq 4000 (single read 50) or a NovaSeq 6000 (SP 100bp flow cell) instrument.

### ChIPmentation

ChIPmentation was performed as previously described (Schmidl et al., 2015). For KLF4 ChIP-seq experiments, 1×10^6^ basal and suprabasal SCC cells were isolated by flow cytometry as described in Figure S4C. H3K27Ac ChIP-seq experiments were conducted with 2×10^6^ cultured TPCs. Cells were fixed, first with ChIP crosslink Gold (Diagenode C01019027) and then with formaldehyde and resuspended in 200 µL of sonication buffer (10mM Tris pH8, 0.25% SDS, 2mM EDTA) supplemented with PIC, and sonicated as described above. Chromatin was then diluted 1:1.5 with equilibration buffer (10mM Tris, 233mM NaCl, 1.66 Triton X-100. 0.166 DOC, 1mM EDTA, PIC) and incubated over night with 4 ug of anti-KLF4 or H3K27Ac antibodies, followed with a 3-hour incubation with 10 µl protein G Dynadeabs (previously blocked in 1% BSA). Immunocomplexes were washed twice with RIPA-LS, RIPA-HS, RIPA-LiCl and 10mM Tris pH8 supplemented with PIC (LS: 10mM Tris-HCl pH8, 140mM NaCl, 1mM pH8 EDTA, 0.1% SDS, 0.1% Na.Deoxycolate, 1% Triton X-100;HS: 10mM Tris-HCL pH8, 1mM EDTA, 500mM NaCl, 1% Triton X-100, 0.1% SDS, 0.1% DOC; LiCl: 10mM Tris-HCl pH8, 1mM EDTA,250mM, 0,5% NP-40, 0.5% DOC), before they were incubated with the tagmentation reaction, using 1 µL Tn5 in 1x Tagmentation buffer (50mM Tris pH8, 25mM MgCl2, 50% v/v dimethylformamide) for 1min at 37 C. Tagmented DNA was washed twice with RIPA-LS and TE, eluted, and crosslinks were reversed overnight in elution buffer (10mM Tris-HCl pH8, 05mM EDTA pH8, 300mM NaCl, 0,4%SDS) with proteinase K. DNA was then purified using QIAGEN Min Elute PCR purification kit (28004) in a final volume of 10 µl. To reduce GC and size bias during the PCR amplification step, the number of PCR cycles were optimized by qPCR and amplification was stopped before saturation, at a cycle number corresponding to a quarter of the maximum fluorescent intensity. Libraries were cleaned and size selected using SPRI beads (AMPureXP beads, Beckman). Multiplexed libraries were sequenced on an IlluminaHiSeq 4000 Genome Analyzer using the 50-base single-end read method.

### RNA-seq library preparation and sequencing

Total RNA was isolated from 3×10^5^ HRas^G12V^, HRas^G12V^;FSOX2^WT^, HRas^G12V^;FSOX2^AA^, and HRas^G12V^;FSOX2^DD^ or SOX2^KO^;FSOX2^AA^ and SOX2^KO^;FSOX2^DD^ GFP-lineage marked SCC cells using Qiazol (Qiagen) and Direct-zol RNA MiniPrep Kits (Zymo Research, R2052). RNA quality was assessed using an Agilent 2100 Bioanalyzer before preparing ribo-depleted, multiplexed, paired end libraries with the Illumina TruSeq Stranded Ribozero Gold kit. Multiplexed libraries were sequenced on an Illumina NovaSeq 6000 using the S1 100bp flow cell.

### 4C library preparation and sequencing

SCC cultures were process and analyzed as described (Rocha et al., 2016; Simonis et al., 2006) with some modifications. 10 million SCC cells were fixed in formaldehyde 2% 10 minutes at RT rocking. After formaldehyde neutralization with glycine 0.125M, cells were washed with cold PBS, resuspended in lysis buffer (50mM Tris, 150mM NaCl, 5mM EDTC, 0.5% NP-40, 1% TX-100, with complete protein inhibitors Roche) and dounce-homogenized to extract nuclei. Nuclei were digested with DpnII overnight (NEB R0543M), followed by an overnight ligation with T4 DNA ligase (Invitrogen 15224). Ligated DNA was reverse-crosslinked overnight with proteinase K and RNAse A and was purified by phenol-chloroform extraction. DNA was digested overnight with the secondary enzyme Csp6I (Fermentas ER0211) and ligated overnight with T4 DNA ligase (NEB M0202). Next, DNA was phenol-chloroform isolated and column purified (GE28-9034-71). Digestion and ligation efficiencies were confirmed after each step by reverse crosslinking, phenol-chloroform DNA extraction and electrophoresis in agarose gel.

Libraries were made using the Expand Long Template PCR system (Roche) with primers designed to target the KLF4 promoter. Primers included Illumina adaptor sequence and a specific barcode for each replicate. Final libraries were purified using columns (GE28-9034-71), size selected with AMPure XP beads (Beckman) and quality assessment on the TapeStation. Multiplexed libraries were sequenced with a NextSeq 500 using the High Output Kit v2.5 (150 Cycles) and paired-end settings.

Demultiplexed reads were mapped to a reduced mm10 genome containing 44bp flanking regions of DpnII restriction sites and interactions were analyzed with 4C-ker to identify cis-interactions as described (Raviram et al., 2016).

### Quantification and statistical analysis

All experiments were carried out single blinded. All in vivo quantitative data were collected from experiments performed in at least triplicate independent tumors and expressed as mean ± s.e.m. Box plots show median, 25^th^ and 75^th^ percentile, minimum, and maximum. Differences between groups were tested using unpaired two-tailed Student’s t-tests, Log-Rank (Mantel Cox) tests, or otherwise indicated in figure legends. Statistical analyses were performed with Graphpad Prism 8 software. Differences were considered significant if p<0.05 and p-values are shown on main and supplemental figures. For clonal competition assays, total population size was counted as percentage of GFP^+^ or RFP^+^ events out of the total live tumor singlets population and expressed as a log_2_ fold enrichment compared to pre-injection values. For KLF4 and FLG2 staining quantification, total signal intensity in the FLG2 channel was divided by the number of cells in each field to generate an average signal intensity per cell per field. For KLF4 basal and suprabasal quantification, basal cells were defined as cells directly adjacent to the integrin α6+ basement membrane. Suprabasal SCC were identified by excluding basal SCC cells and stromal cells. KLF4 staining intensity was then measured and divided by either the number of basal or suprabasal cells to get KLF4 staining intensity per cell per field.

Two approaches were taken for gating strategies in the α6- and β1-integrin high vs low comparison. To normalize for sample-to-sample variation in the HRas^G12V^ model, ɑ6^hi^/β1^hi^ and ɑ6^lo^/β1^lo^ gates in the GFP^+^ SCC cell lineage were defined relative to α6- and β1-integrin levels of GFP^-^ skin epithelial cells. For the Sox2^KO^ model, we ensured equal spectral compensation by matching compensation bead (BD #552844 and #552843) intensity within a narrow range for every run. FACS gates were defined on one tumor and applied to every other tumor in the cohort.

Statistical significance for gene set intersections was calculated using the sizes of the two gene sets, the overlap between the two gene sets, and the size of all genes in the background mouse genome, using online calculation tools (http://nemates.org/MA/progs/overlap_stats.html).

### ChIP-seq analysis

Two biological replicates were sequenced in each ChIP-seq experiment. Mouse SOX2 ChIP-seq and human spike-in ChIP-seq reads were aligned to the mm10 mouse genome or hg38 human genome using bowtie2, version 2.3.4.1 (Langmead and Salzberg, 2012). Aligned reads with a quality score of <30 were discarded with Samtools, version 1.9 (Li et al., 2009) and PCR duplicates were discarded using Picard-tools, version 1.88 (http://broadinstitute.github.io/picard/). IGV viewer files were generated using igvtools, version 2.4.1 (Robinson et al., 2011) with the following settings: -z 5 -w 25 -e 250. A scale factor was calculated from the number human spike-in reads aligned to the human genome. The sample with the lowest human reads was set as 1 and the rest of the samples scaled down to match it using picard-tools downsampleSam, STRATEGY=Chained, RANDOM_SEED=8 to randomly remove reads from the .bam files with a probability equal to the scale factor. ChIP-seq peaks were called using MACS2, version 2.1.1 (Feng et al., 2012; Zhang et al., 2008) in comparison to input controls with default settings and a -q .01 (Sox2^KO^) or -q 0.001 (HRas^G12V^) cutoff. Motif discovery was performed using MEME-ChIP, version 5.0.2 (Bailey et al., 2009; Machanick and Bailey, 2011) on the ChIP peak summits ±100bp with default settings. Peaks containing the SOX motif were combined to generate a “peakome” that comprises all called peaks from all samples of an experiment. Motif containing peak percentages were calculated based on the number of motif containing peaks within all peaks in the peakome. The Genomic Regions Enrichment of Annotations Tool (GREAT, (McLean et al., 2010)) was used to determine the SOX2 peak distributions (from nearest TSS) and identify nearest neighbor genes relative to SOX2 bound sites. Two replicates were sequenced for KLF4 and H3K27Ac, and one replicate was sequenced for MED12. Analyses were performed as described for SOX2 ChIP-seq with some modifications. MACS was used with a -q 0.05 cutoff for KLF4 ChIP-seq studies, and the –broad option with the default q cutoff was used for H3K27Ac ChIP-seq studies. No spike-in normalization controls were used in these experiments.

To compare overall ChIP-seq signals and distributions, Deeptools, version.3.2.1 (Ramírez et al., 2014) was used to create a .bw file from the .bam files (--exactScaling, RPGC normalization, and bin size 10 settings), quantifying the signal using computeMatrix (reference-point mode, -b 500 and -a 1500 settings). Heatmaps, profiles (histograms), and distributions were generated from the matrix file.

RNA-seq and ChIP-seq data were integrated by extracting the closest up and downstream genes associated with each peak using GREAT and overlapping gene lists. Gene ontology analyses were performed using AmiGO, version 2.5.12 (Carbon et al., 2009).

For statistical analysis, reads within all peaks in the peakome genomic regions were counted from down-sampled .bam files (SOX2) or original .bam files (KLF4) using HTSeq (Anders et al., 2015) and differentially bound regions were determined using DESeq2 (Love et al., 2014) with a p<0.1 (KLF4 ChIP-seqs), p<0.05 and log_2_ fold change cutoff of 0.585 (1.5 fold change) (HRas^G12V^; SOX2 and SOX2^KO^; SOX2 ChIP-seqs). Regions with low coverage (<10 read counts) were discarded.

### RNA-seq analysis

Three replicates were sequenced for each RNA-seq. RNA-seq data were analyzed using the Tuxedo Suite of tools (Kim et al., 2013; Langmead and Salzberg, 2012; Trapnell et al., 2012) RNA-seq reads were aligned to the mouse mm10 genome using the tophat/bowtie2 aligner (tophat v2.1.1, bowtie v2.3.4.1) and –no-coverage-search option. Transcriptome quantification and differential expression (DE) analysis was performed using the Cufflinks/Cuffquant/Cuffdiff/CummeRbund pipeline (cufflinks v2.2.1) with default parameters. DE analysis was performed on HRas^G12V^;FSOX2^AA vs DD^ and SOX2^KO^;FSOX2^AA vs DD^ conditions. Area proportional diagrams were generated with BioVenn (Hulsen et al., 2008).

## Notes

### Competing Interest Statement

The authors have declared no competing interest.

