## Supplementary figures and images for "SOX2-phosphorylation toggles a bistable differentiation-switch in squamous cell carcinoma"

### Figure S1

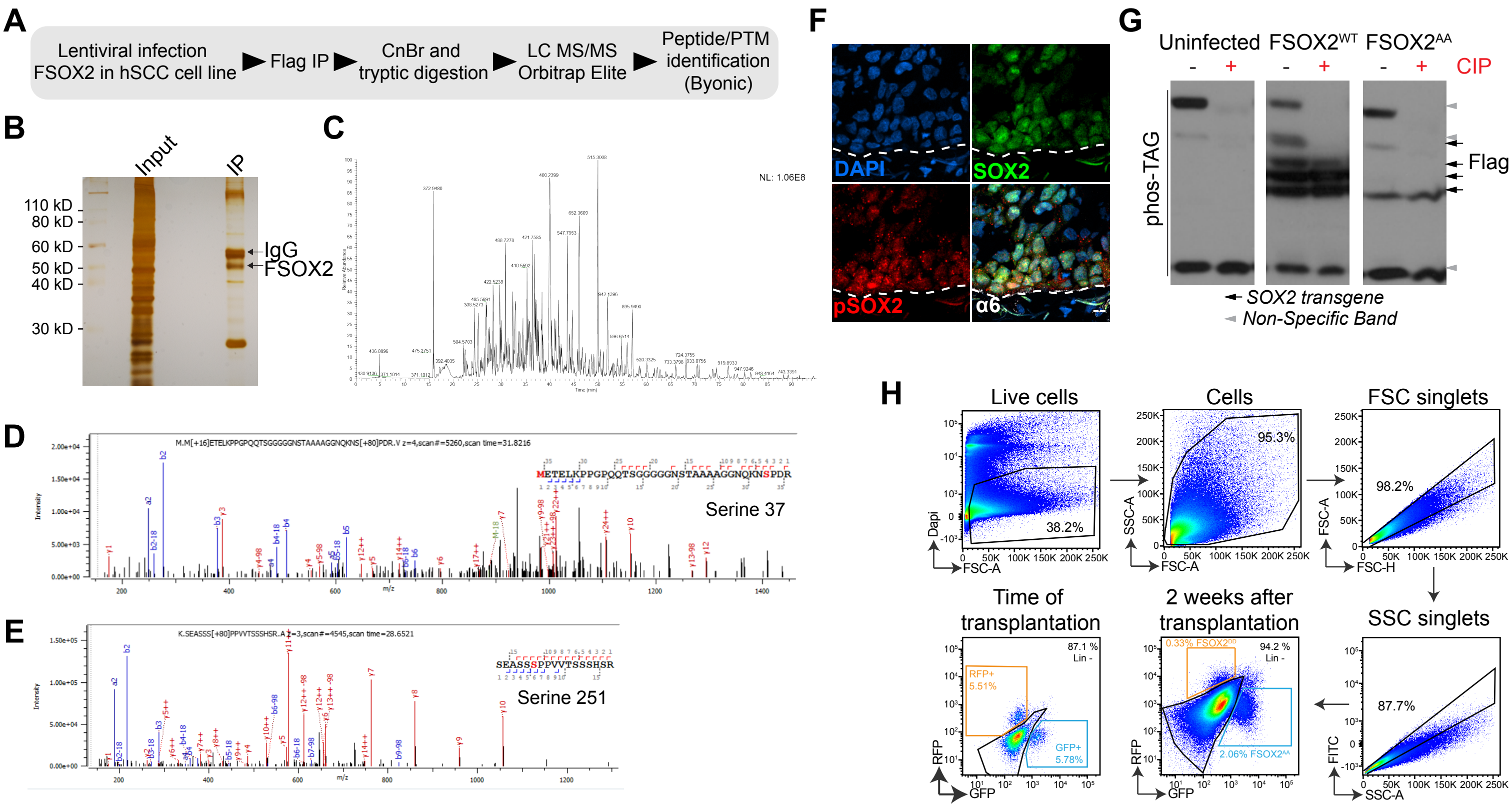

### Figure S2

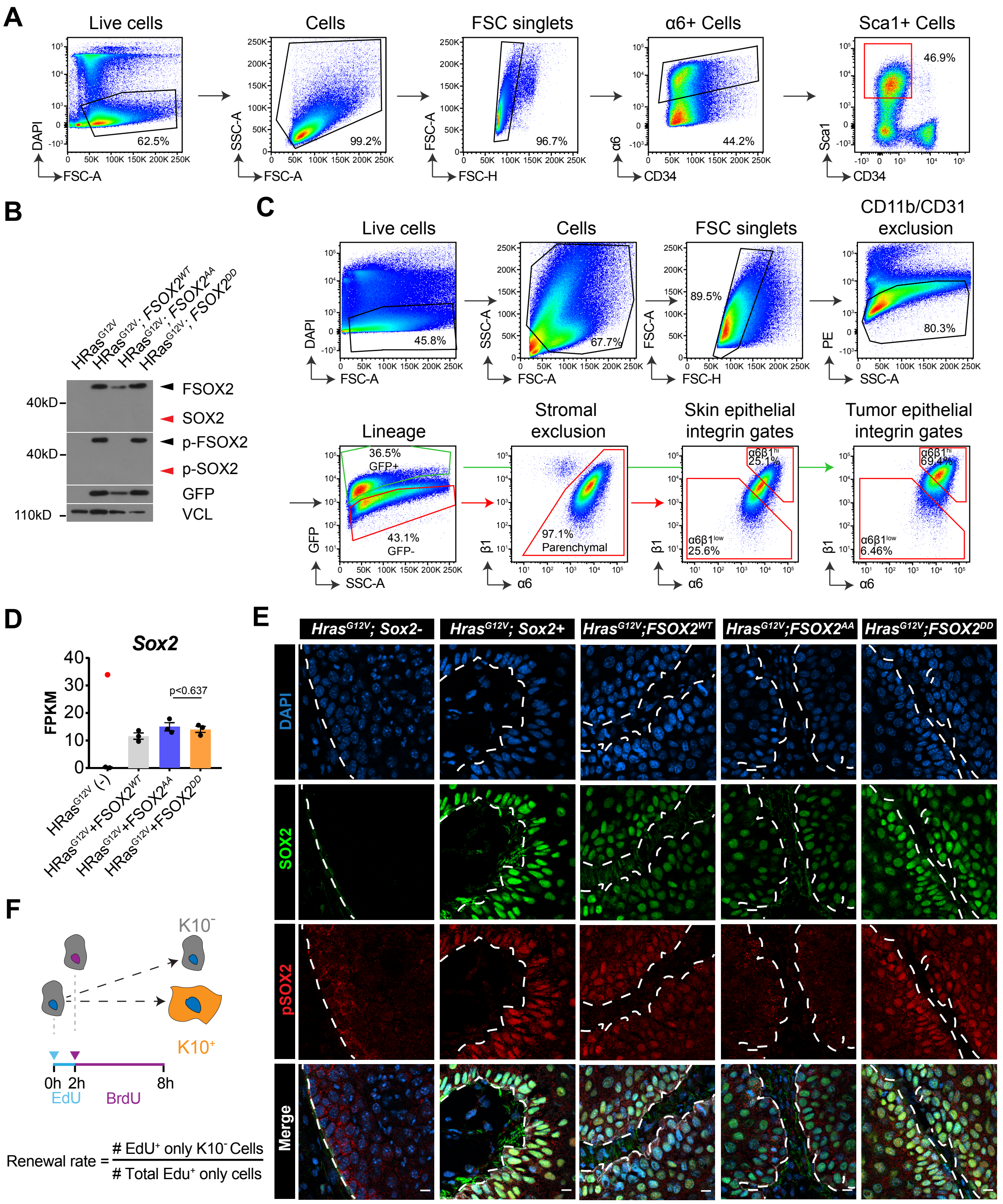

### Figure S3

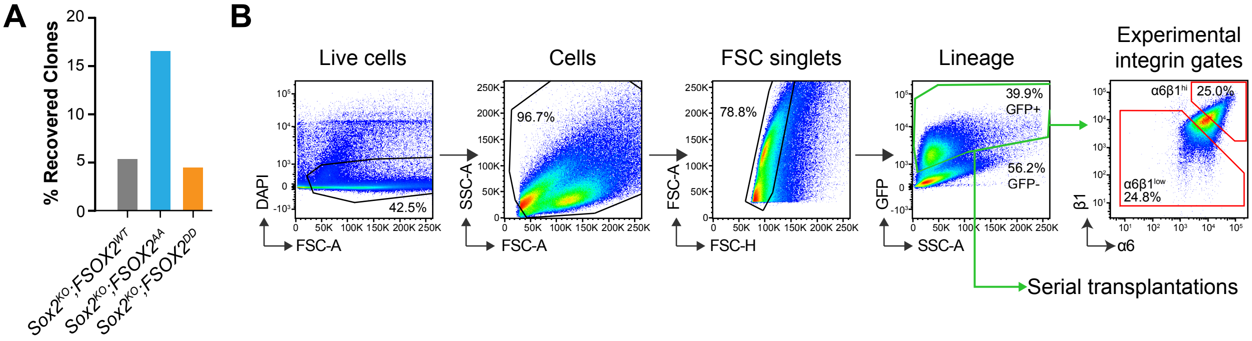

### Figure S4

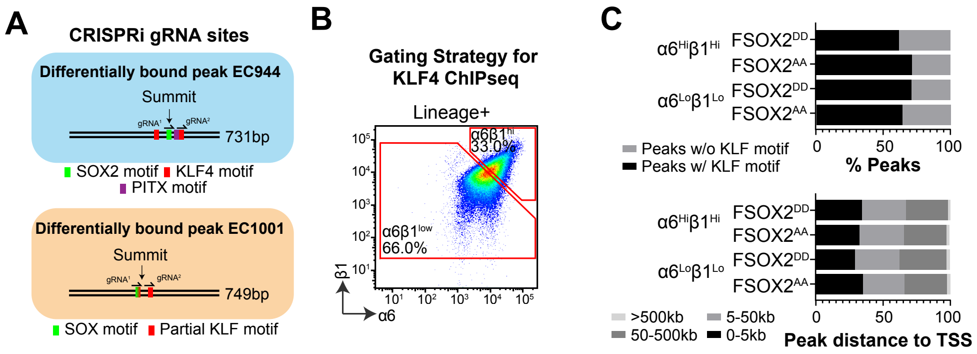
